# OrthoSimulator: Empirically trained simulations for ground-truth benchmarking of orthology inference

**DOI:** 10.64898/2026.11.12.766176

**Authors:** Jonathan Holmes, Steven B. Kelly, Laurence J. Belcher

## Abstract

Orthology inference is foundational for comparative genomics, enabling the reconstruction of phylogenetic relationships and transfer of biological knowledge between species. Methods often infer different orthogroups and orthologous relationships, making robust benchmarking essential for evaluating their accuracy. Existing resources such as the Quest for Orthologs benchmark service and OrthoBench provide valuable benchmarks grounded in real biological data and expert curation. However, the true evolutionary histories in all of these benchmarks are unknown, and therefore the true accuracy of orthology inference methods is unknown. Simulations provide an approach to evaluate accuracy where the complete ground truth is known. However, simulations are only useful for benchmarking if they realistically reflect empirical data. Here, we introduce OrthoSimulator, a Python package for generating realistic, empirically calibrated benchmark datasets with known orthogroups, orthologous relationships, gene trees, and evolutionary events. We demonstrate its utility by benchmarking widely used orthology inference methods across simulations calibrated from multiple methods and empirical datasets. We show that method-to-method variability in ortholog inference accuracy is robust to variation in simulation parameters. We further provide a detailed evaluation of ortholog inference accuracy through independent cross-validation, and show that OrthoFinder is the most accurate of the four methods tested. OrthoSimulator is freely available at https://github.com/lauriebelch/OrthoSimulator.

**Significance Statement:** Comparative genomics relies on orthology, the identification of genes that descend from a single gene in a common ancestor. Many methods infer orthologs, and they often disagree, but their true accuracy is hard to measure because the true history of real genes is never known. We present OrthoSimulator, which learns from real genomes to simulate realistic gene families whose full history is recorded. The simulations complement existing benchmarks built from real genes by adding complete ground truth. We use them to compare four widely used methods and to measure errors that cannot be seen in real data, and the simulated genomes can test any method that reconstructs gene family evolution.

## Introduction

Orthology inference is fundamental to biological research, providing the foundation for comparative genomics and phylogenetics and enabling the transfer of biological knowledge between species. The reliability of downstream biological analyses depends on accurate orthology inference. Benchmarking these methods is therefore essential for assessing this accuracy, ideally by comparing inferred orthogroups and orthology relationships against datasets for which the true relationships are known. Since the concept of orthology was formalised over fifty years ago (Fitch 1970) many inference methods have been developed, each making different methodological choices and assumptions (Li et al. 2003; Kaduk and Sonnhammer 2017; Derelle et al. 2020; Linard et al. 2021; Cosentino et al. 2024; Majidian, Nevers, et al. 2025; Emms et al. 2026). These differences in approach lead to differences in orthogroups and orthologs between methods, and it can be unclear whether these differences correspond to genuine differences in accuracy. This is important because errors in orthology inference can propagate into downstream analyses, leading to inaccurate phylogenetic trees, mistaken inferences in gene function, and confusion in our understanding of the evolutionary history and diversity of life on earth. Robust benchmarking is therefore essential for evaluating orthology inference methods and ultimately improving their accuracy and reliability (Gabaldón et al. 2009).

The orthology inference community has developed invaluable empirical benchmarking resources for this purpose. The Quest for Orthologs (QfO) benchmark service provides a widely used framework for comparing orthology methods across a shared set of 78 reference proteomes spanning Bacteria, Archaea, and Eukaryotes (Gabaldón et al. 2009; Altenhoff et al. 2016; Linard et al. 2021; Nevers et al. 2022). QfO provides multiple benchmark types, including expert-curated ortholog sets such as VGNC (Jones et al. 2023) and TreeFam (Schreiber et al. 2014), as well as species-tree discordance tests. OrthoBench provides a complementary benchmark based on 12 metazoan species, focusing on orthogroup inference using expert-curated reference orthogroups (Trachana et al. 2011; Emms and Kelly 2020). These resources are extremely valuable because they are grounded in real biological data and expert knowledge. However, empirical benchmarks necessarily have limitations. Expert curation is labour-intensive, meaning that reference datasets are limited in size and scope. For example, OrthoBench contains 70 reference orthogroups across the twelve species, representing only 0.75% of all genes across those genomes. More fundamentally, the true evolutionary history of empirical data is unknown, meaning that even carefully curated reference sets remain inferred rather than observed and may contain errors. Empirical benchmarks therefore cannot provide complete ground truth, which would require knowing the true orthogroup membership and pairwise orthologous relationships of all genes.

Constructing representative benchmarks is complicated by the diversity of evolutionary processes that shape orthogroups. Each orthogroup, defined as a set of genes descended from a single gene in the last common ancestor of a group of species, has arisen through a distinct evolutionary history. Orthogroups do not always evolve through simple patterns of speciation, duplication, and loss within the neat confines of a species tree (Altenhoff et al. 2019; Emms and Kelly 2020). Gene trees can also differ from the species tree because of incomplete lineage sorting (ILS), creating discordance even in the absence of gene duplication and loss (Degnan and Rosenberg 2009). Some evolve rapidly, whereas others remain highly conserved. Genes can also fuse together or split apart, and shared domains or motifs can create misleading patterns of sequence similarity (Durrens et al. 2008; Kristensen et al. 2011). These complications can produce hidden paralogy, erroneously split or merged orthogroups, and ambiguous similarities that mislead methods (Natsidis et al. 2021). They can also cause some orthology relationships to be missed entirely and can lead to genes being erroneously left out of orthogroups. Different tools handle these phylogenetic challenges in different ways, and it is thought that these differences can cause trade-offs between precision and recall (Altenhoff et al. 2016; Glover et al. 2019; Nevers et al. 2022). Precisely due to these complexities, creating large and reliable datasets manually is impractical. Manually created benchmarks can therefore be limited in the range and scale of evolutionary processes that are represented.

Simulations can provide a complementary approach to the issues surrounding benchmarking. The precise evolutionary history of every orthogroup, including every gene gain, duplication, and loss, can be recorded directly, allowing inferred orthogroups and orthologous relationships to be compared against their true values. Simulations can also be scaled, replicated, and modified to test performance under different evolutionary scenarios. Previous simulation studies have demonstrated the value of this approach for understanding orthology inference errors. For example, variation in rates of gene duplication and loss can alter the balance between recall and precision across different methods (Dalquen et al. 2013). Similarly, variation in gene evolutionary rate can cause detection failure, generating false presence/absence patterns that confound downstream analyses (Natsidis et al. 2021). Although these studies identified specific causes and consequences of orthology inference errors, they were designed to test particular evolutionary processes rather than to generate general-purpose benchmark datasets.

The ability to simulate gene trees and multiple sequence alignments at scale makes it possible to generate large orthogroup datasets for which the complete evolutionary history is known. This offers an important advantage over curated empirical reference datasets, which can offer limited information about the underlying evolutionary events that cause errors to arise. With simulations all evolutionary events can be recorded and a complete evolutionary history is known, making it possible to distinguish the circumstances responsible for inference failure. This complete evolutionary ground truth allows method developers to identify the evolutionary events associated with inference errors and test whether methodological changes resolve them. However, for these important insights to be realised simulated benchmarking must be able to reproduce the properties and evolution of real orthogroups.

Here, we present OrthoSimulator, a Python package for generating realistic orthogroup benchmark datasets with complete evolutionary ground truth. OrthoSimulator uses empirical orthogroups inferred by any orthology inference method to estimate the evolutionary parameters needed to generate new simulated datasets that reflect the properties of real data. We demonstrate OrthoSimulator using multiple empirical datasets and orthology inference methods, and use the resulting simulations to benchmark orthogroup and ortholog inference against known ground truth. By calibrating simulations independently from the outputs of different inference methods, we further test whether benchmarking results are robust to the empirical dataset used to generate them. The complete evolutionary history also enables evaluation of otherwise difficult-to-measure errors, such as hidden paralogy, and testing under evolutionary complications including incomplete lineage sorting. OrthoSimulator therefore provides a scalable, empirically grounded complement to existing reference benchmarks, allowing orthology methods to be evaluated across complete datasets for which the true orthogroups, gene histories, and orthology relationships are known.

## Results

### The OrthoSimulator workflow

OrthoSimulator is a Python package for generating empirically calibrated orthogroup benchmark datasets with complete evolutionary ground truth. It takes as input a set of orthogroups inferred by any orthology inference method, including OrthoFinder (Emms et al. 2026), FastOMA (Majidian, Nevers, et al. 2025), SonicParanoid2 (Cosentino et al. 2024), Broccoli (Derelle et al. 2020), or any other tool that provides orthogroups in OrthoXML format (Schmitt et al. 2011; Yazdizadeh Kharrazi et al. 2025). OrthoSimulator also requires the proteomes that were used to infer the orthogroups and a rooted, ultrametric species tree but can read these directly from the output of many orthology inference tools.

OrthoSimulator then proceeds through three major steps. First, it extracts the sequences from each orthogroup and estimates parameters that can be measured directly from the empirical data, including sequence-evolution parameters, branch-rate variation, alignment properties, orthogroup size, and presence or absence across species. Second, it uses a genetic algorithm to optimise the duplication rate, loss rate and orthogroup-birth parameters that cannot be measured directly from orthogroup assignments. Third, OrthoSimulator uses these parameters to simulate a new set of orthogroups. The final output is a complete simulated benchmark dataset containing proteomes, alignments, gene trees, orthogroup membership tables and pairwise ortholog relationships, together with a per-orthogroup log file recording every event and parameter value sampled during simulation (Figure 1).

**Figure 1:**
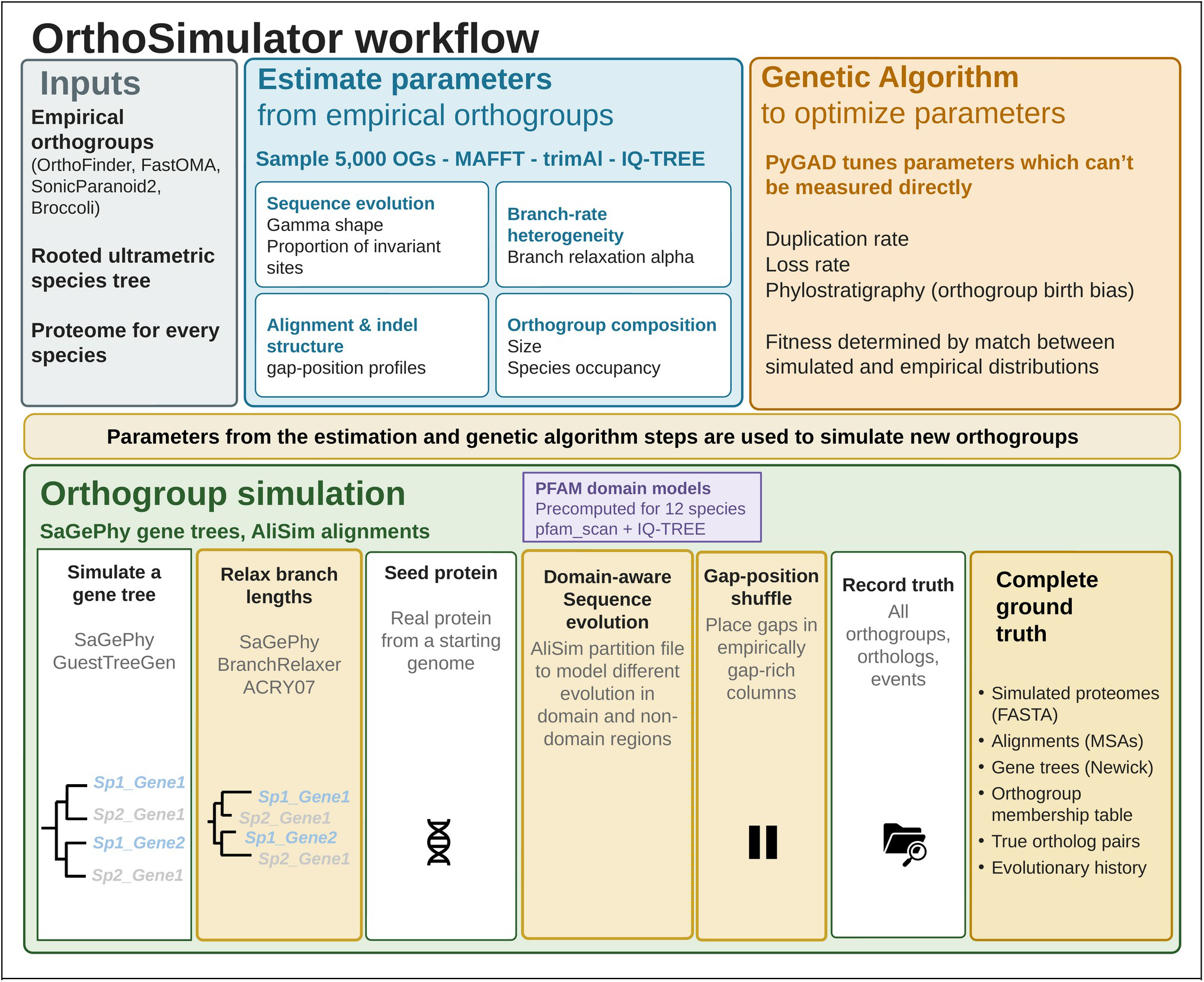
OrthoSimulator workflow. OrthoSimulator takes three inputs: a set of empirical orthogroups inferred by any orthology method, a rooted ultrametric species tree, and a proteome for every species. From these, OrthoSimulator estimates parameters in two ways. It measures sequence-evolution parameters, branch-rate heterogeneity, alignment and indel structure, and orthogroup composition directly from the empirical orthogroups. It then uses a genetic algorithm to tune the parameters that cannot be measured directly by comparing the distributions produced by trial simulations against the empirical distributions. These trained parameters are then used to simulate new orthogroups. For each orthogroup, OrthoSimulator simulates a gene tree and relaxes its branches with SaGePhy and seeds it with a real protein sequence from a starting genome. It then simulates sequence evolution with IQ-TREE’s AliSim, using a partition file that lets PFAM domain and non-domain regions of the seed protein evolve under different models (drawing on PFAM domain models precomputed for a set of model species, or computed by the user for their own proteome), and repositions simulated gaps to match empirically gap-rich alignment columns. OrthoSimulator records the complete evolutionary history of every simulated orthogroup. The output is a full simulated dataset: proteomes, alignments, gene trees, an orthogroup membership table, true ortholog pairs, and the underlying evolutionary history of every simulated event.

OrthoSimulator seeds each simulated orthogroup with a real protein sequence drawn from an empirical proteome. Where the seed protein carries a PFAM domain (Sonnhammer et al. 1997), OrthoSimulator evolves that region of the simulated alignment under a domain-specific substitution model, so that domain and non-domain sequence evolve under different constraints. We provide precomputed PFAM domain models for twelve species (Homo sapiens, Arabidopsis thaliana, Saccharomyces cerevisiae, Escherichia coli, Mnemiopsis leidyi, Danio rerio, Gallus gallus, Mus musculus, Chlamydomonas reinhardtii, Drosophila melanogaster, Triticum aestivum, and Haloferax volcanii), and a module that builds these models for any other proteome the user supplies (see ‘PFAM domain model extraction’ in Methods).

For the principal benchmarking analyses, we use the baseline duplication-loss model, in which gene lineages coalesce at speciation and ILS is therefore not introduced. OrthoSimulator additionally provides an optional multispecies-coalescent extension for introducing ILS using a single parameter.

### OrthoSimulator generates realistic benchmark datasets

OrthoSimulator can generate benchmark datasets with or without incomplete lineage sorting (ILS). We first assess simulation realism and benchmark orthology inference without ILS, then introduce ILS to examine its effects on inference accuracy. To provide informative benchmarks, simulated orthogroups should reproduce key properties of empirical data. We therefore generated eight simulated datasets, each containing 10,000 orthogroups, calibrated from the outputs of four orthology inference methods (OrthoFinder, FastOMA, SonicParanoid2 and Broccoli) on QfO and OrthoBench. We compared eight properties of orthogroup composition, sequence evolution, alignment structure and gene tree shape between the simulated and corresponding empirical datasets (Methods). The SonicParanoid2-calibrated QfO dataset is shown in Figure 2, with the remaining comparisons shown in Supplementary Figures 1–7.

**Figure 2:**
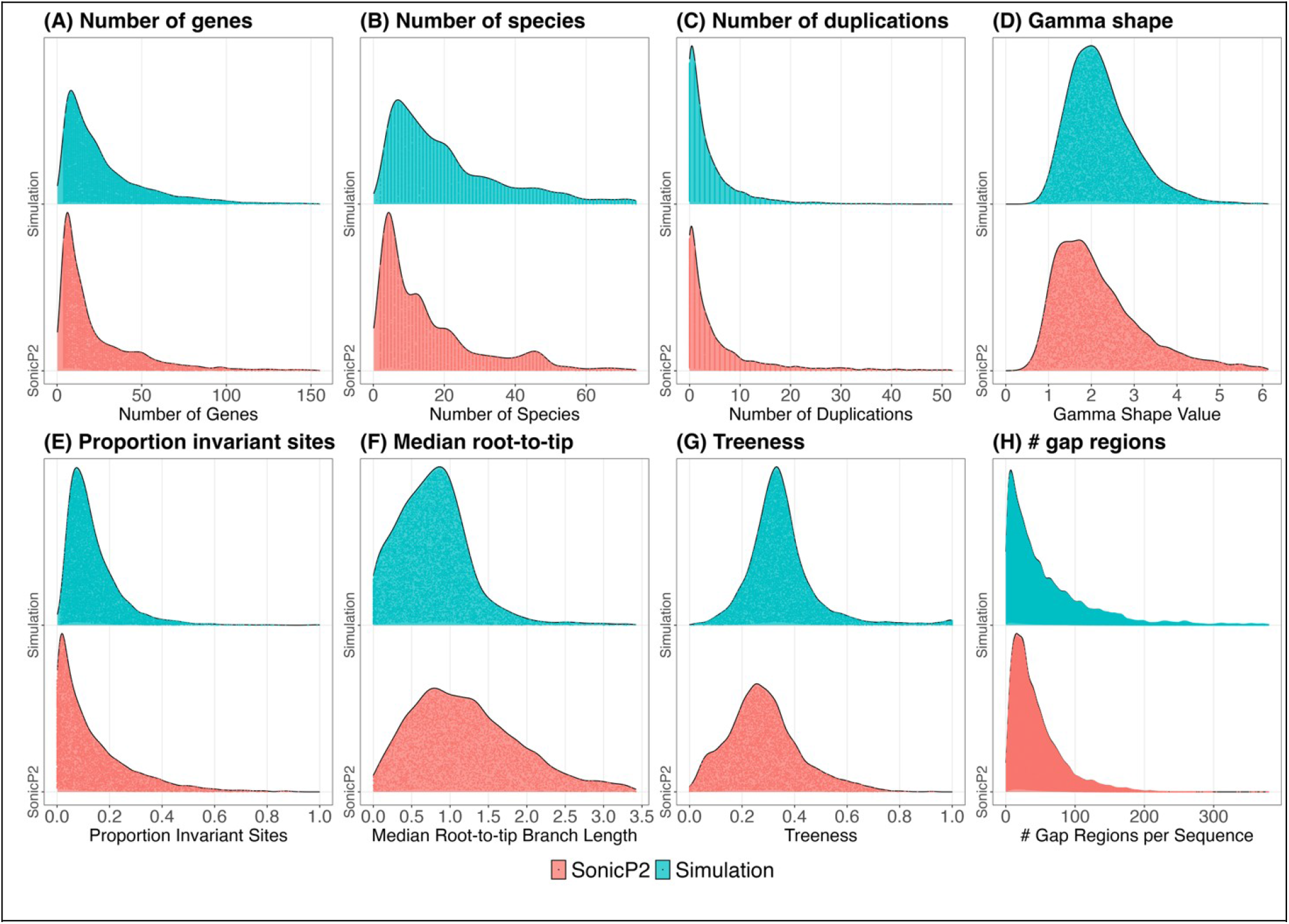
OrthoSimulator reproduces the properties of empirical orthogroups. Distributions of eight orthogroup and gene-tree properties for the SonicParanoid2-trained QfO simulation (teal) compared against the empirical SonicParanoid2 orthogroups used to train it (red). (A) Number of genes per orthogroup. (B) Number of species represented per orthogroup. (C) Number of duplications per orthogroup. (D) Gamma shape parameter (alpha) of the among-site rate-variation model. (E) Proportion of invariant sites per alignment. (F) Median root-to-tip distance per gene tree, a measure of evolutionary rate. (G) Treeness, the proportion of total branch length found on internal branches. (H) Number of gap regions per aligned sequence. Equivalent comparisons for the other three training methods (OrthoFinder, Broccoli, FastOMA) and for the OrthoBench dataset are shown in Supplementary Figures 1-7. In each panel, the vertical axis shows relative density (arbitrary units, comparable within a panel but not between panels).

The simulated orthogroups closely matched the empirical data for several properties relevant to orthology inference. For orthogroup size, the difference between the simulated and empirical distributions was only 35% of the largest difference observed between two empirical methods analysing the same genomes (Figure 2A). For the number of species represented per orthogroup, this value was 88% (Figure 2B), and for the number of duplications per orthogroup it was 59% (Figure 2C). Thus, for each of these properties, the simulated dataset in Figure 2 was more similar to the empirical data used for calibration than the most divergent pair of empirical orthology methods were to one another. The true number of historical gene losses cannot be observed directly in empirical data, and therefore loss rate could not be validated in the same way. However, the genetic algorithm jointly estimates duplication rate, loss rate, and birth bias to reproduce measurable properties including orthogroup size, species representation, and duplication counts. The estimated loss rate is therefore constrained by these empirical properties, although it cannot be uniquely identified from them.

Evolutionary rates were also reproduced closely. For median root-to-tip distance in gene trees, the difference between the simulated and empirical distributions was 90% of the largest difference between empirical methods (Figure 2F). The sequence-evolution parameters contributing to these patterns were similarly well reproduced, with corresponding values of 95% for the alpha parameter of the gamma distribution governing among-site rate variation (Figure 2D) and 50% for the proportion of invariant sites per alignment (Figure 2E).

Two properties were reproduced less closely. For gene-tree treeness, the difference between the simulated and empirical distributions was 1.9 times the largest difference between empirical methods (Figure 2G), while for the number of gap regions per alignment it was 1.7 times larger (Figure 2H). Nevertheless, both showed similar overall distributions between simulated and empirical data, consistent with the branch-length and indel-simulation simplifications discussed below. Overall, OrthoSimulator reproduced the major empirical properties most relevant to orthology inference, providing simulated datasets suitable for ground-truth benchmarking.

### Orthogroup benchmarking resolves distinct inference errors

We benchmarked all four orthology inference methods on each of the eight simulated datasets. The complete ground truth allows inference accuracy and distinct error profiles to be assessed across thousands of orthogroups, extending benchmarking beyond the limited reference sets available from empirical curation. We evaluated orthogroup inference using seven complementary measures that capture different types of error: recall, the proportion of genes from each reference orthogroup recovered in predicted orthogroups; precision, the proportion of genes in predicted orthogroups belonging to the corresponding reference orthogroup; fission, the splitting of reference orthogroups across multiple predicted orthogroups; fusion, the merging of genes from different reference orthogroups; missing genes; missing reference orthogroups; and entropy, which measures fragmentation of reference orthogroups across predicted orthogroups. Where measures were calculated for individual reference orthogroups, they were combined across the dataset using orthogroup-size weighting (Emms et al. 2026). We additionally summarised performance across all seven measures using a composite z-score, where higher is better. Results for the QfO simulations are shown in Figure 3 (see also Supplementary Figure S1.1 for the OrthoBench results).

**Figure 3:**
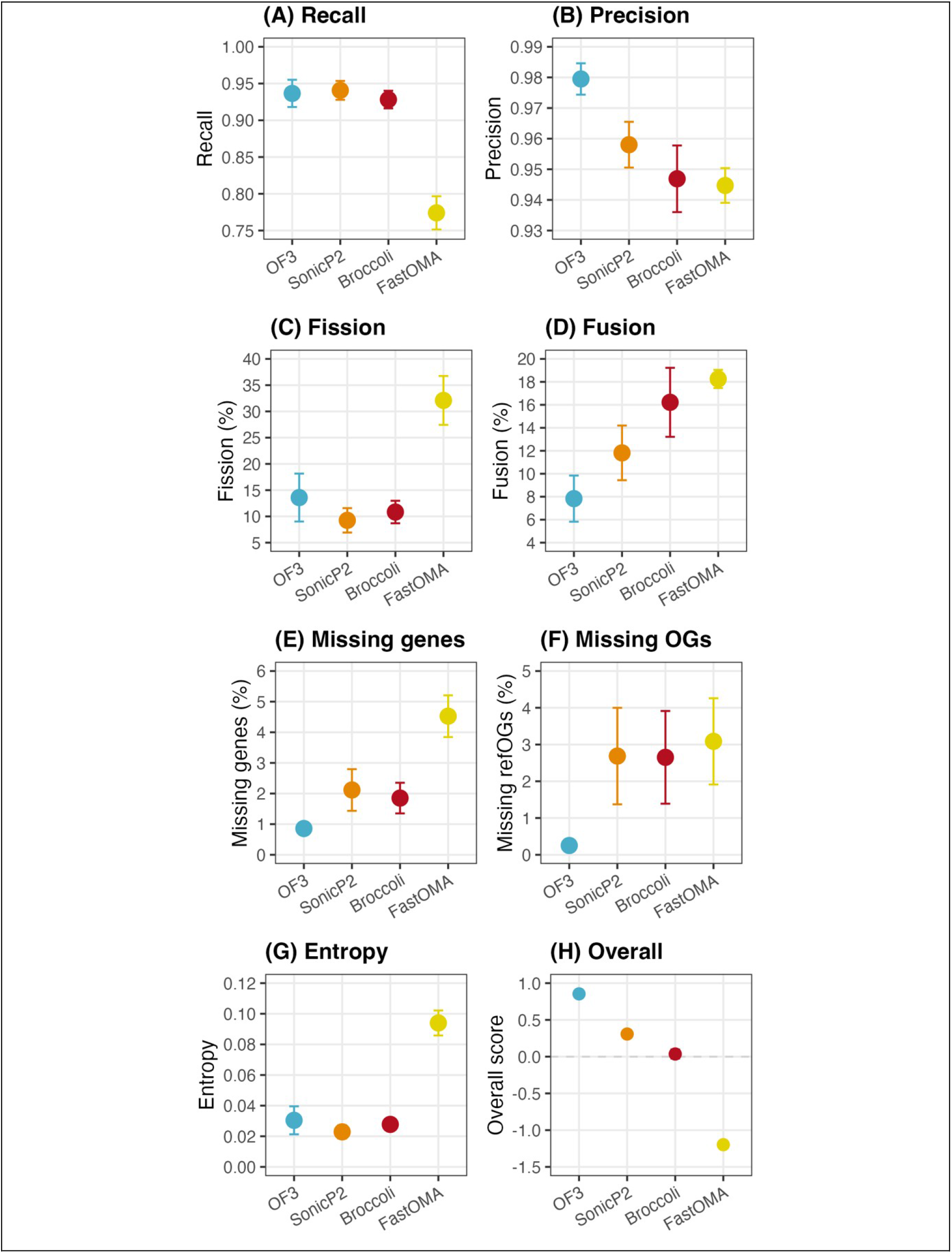
Orthogroup-level benchmarking of four orthology inference methods on the QfO-trained simulations. Each panel shows one benchmark measure for OrthoFinder (OF3), SonicParanoid2 (SonicP2), Broccoli, and FastOMA. (A) Recall: the proportion of genes in each reference orthogroup recovered in the predicted orthogroups. (B) Precision: the proportion of genes in each predicted orthogroup that belong to the corresponding reference orthogroup. (C) Fission: the percentage of reference orthogroups split across more than one predicted orthogroup. (D) Fusion: the percentage of reference orthogroups whose genes were merged with genes from at least one other reference orthogroup within the same predicted orthogroup. (E) Missing genes: the percentage of reference-orthogroup genes absent from every predicted orthogroup. (F) Missing OGs: the percentage of reference orthogroups entirely absent from the predicted set. (G) Entropy: the size-normalised Shannon entropy of each reference orthogroup’s distribution across predicted orthogroups, a measure of fragmentation. (H) Overall: the composite z-score summarising performance across all seven measures in A-G. Equivalent results for OrthoBench are shown in Supplement S1.

OrthoFinder had the highest overall composite score across the QfO simulations (0.85 ± 0.04, mean ± standard error across the four calibrations; Figure 3H), followed by SonicParanoid2 (0.31 ± 0.02), Broccoli (0.04 ± 0.06), and FastOMA (−1.20 ± 0.04). The individual measures also revealed distinct performance profiles between methods. OrthoFinder had the highest precision (0.979 ± 0.005), the lowest rate of orthogroup fusion (7.8 ± 2.0%), and the lowest proportions of missing genes (0.86 ± 0.14%) and missing reference orthogroups (0.25 ± 0.05%). Recall was similar for OrthoFinder (0.937 ± 0.019) and SonicParanoid2 (0.941 ± 0.013). FastOMA showed a different balance of errors, with lower recall (0.774 ± 0.023) and a higher rate of orthogroup fission (32 ± 5%), while fusion occurred in 18.3 ± 0.8% of reference orthogroups. Thus, while OrthoFinder showed the strongest overall performance in this benchmark, the complete ground truth also resolves the distinct strengths and error profiles of each method across multiple aspects of orthogroup inference.

### Ortholog benchmarking directly measures inference accuracy

Empirical benchmarking of pairwise orthology remains limited in both scale and taxonomic coverage. The QfO benchmarking service, for example, includes a benchmark based on curated orthologous relationships from the Vertebrate Gene Nomenclature Committee (VGNC). However, this contains a limited set of reference relationships covering only four of the 78 QfO reference species, all mammals (Nevers et al. 2022). These expert-curated relationships provide a valuable reference, but remain inferences from empirical data rather than a complete record of the true evolutionary relationships. OrthoSimulator provides complete simulation ground truth containing every true pairwise orthologous relationship across all species in each simulated dataset, allowing ortholog inference to be evaluated directly. We classified inferred ortholog pairs as true positives, false positives, or false negatives and calculated precision, recall, and their harmonic mean, the F1-score, for each method (Figure 4; see also Supplementary Figure S1.2).

**Figure 4:**
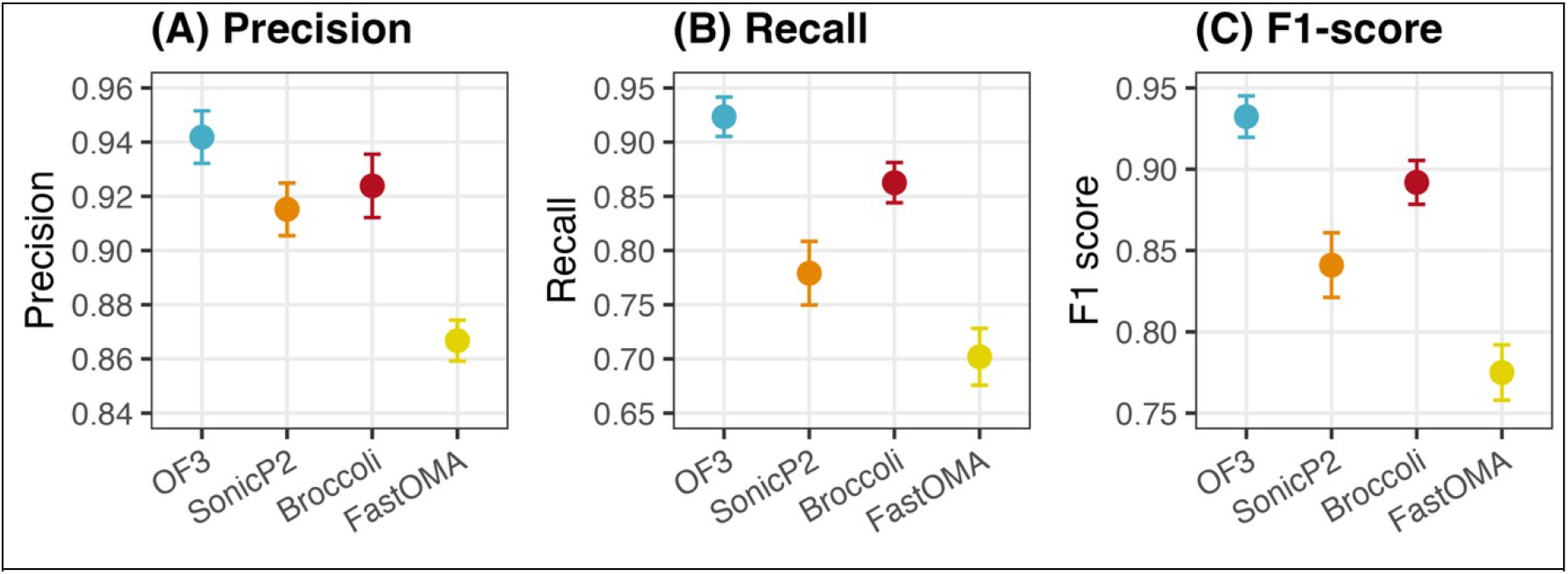
Ortholog-pair benchmarking of four orthology inference methods on the QfO-trained simulations. (A) Precision, (B) recall, and (C) F1-score for pairwise ortholog inference, calculated by comparing each tool’s inferred ortholog pairs against the complete set of true simulated pairs. Equivalent results for OrthoBench are shown in Supplement S1.

OrthoFinder had both the highest precision (0.942 ± 0.010) and highest recall (0.923 ± 0.018), giving the highest F1-score (0.932 ± 0.013). Broccoli had the next highest F1-score (0.892 ± 0.013), followed by SonicParanoid2 (0.841 ± 0.020) and FastOMA (0.775 ± 0.017). The difference between OrthoFinder and the next highest-scoring method was 0.040 in F1-score. These results show that OrthoSimulator can resolve differences in pairwise ortholog inference using direct precision and recall against known evolutionary ground truth. OrthoSimulator therefore provides a reproducible method for benchmarking features of orthology inference at a scale and flexibility that were not previously possible.

### OrthoSimulator enables benchmarking under incomplete lineage sorting

Incomplete lineage sorting (ILS) can generate discordance between gene trees and species trees and may affect orthology inference in some evolutionary settings (Pamilo and Nei 1988; Degnan and Rosenberg 2006). OrthoSimulator provides an optional extension that allows ILS to be introduced explicitly, enabling controlled tests of its effects on orthology inference. We used a no-ILS baseline for the principal benchmarks because the appropriate magnitude of ILS cannot be estimated from the empirical orthogroup data used for calibration.

The extension reshapes simulated gene trees under the multispecies coalescent (Rannala and Yang 2003; Degnan and Salter 2005), while retaining duplication events as fixed, dated ground truth using a bounded coalescent (Rasmussen and Kellis 2012). ILS is controlled by a single parameter, theta, allowing users to vary the extent of coalescent discordance without specifying additional demographic parameters. To demonstrate this capability, we generated an additional set of 10,000 orthogroups using the same calibrated parameters as the OrthoFinder-QfO simulation, but with ILS introduced using theta = 10, and repeated the benchmarking of all four orthology inference methods.

Introducing ILS reduced performance on several measures but retained the broad differences between methods observed in the baseline simulations (Supplementary Figure 10). For example, on the OrthoFinder-calibrated QfO simulation, OrthoFinder recall fell from 0.901 to 0.886 and orthogroup fission rose from 21.1% to 26.8%, while mean ortholog F1 across methods changed little (0.832 without ILS, 0.841 with ILS). Thus, OrthoSimulator can be used to extend ground-truth benchmarking to datasets containing ILS, providing a controlled way to examine how gene tree-species tree discordance affects orthology inference.

### Ground truth allows the benchmarking of hidden paralogy

A particular advantage of simulated data is that the complete evolutionary history enables evaluation of errors that are difficult to identify reliably in empirical datasets. One example is hidden paralogy (an example is shown in Supplementary Figure 8), where genes from different species appear to be orthologous but are in fact paralogs because different copies were retained following an ancestral duplication and differential loss (Doolittle and Brown 1994; Niimura and Nei 2007; Supplementary Figure 8). Because OrthoSimulator records every duplication and loss event, these hidden-paralog pairs can be identified directly from the simulated ground truth.

We quantified a hidden-paralog rejection rate, defined as the proportion of hidden-paralog pairs within the same inferred orthogroup that were correctly not called as orthologs. A value of 1 therefore indicates that all hidden paralogs were correctly distinguished from orthologs, while lower values indicate that a greater proportion were incorrectly called as orthologs. Broccoli had the highest hidden-paralog rejection rate (0.82 ± 0.01), followed closely by OrthoFinder (0.80 ± 0.02), while FastOMA had a lower rejection rate (0.66 ± 0.01) (Figure 5; Supplementary Figure S1.3 for the OrthoBench-calibrated simulations). Under incomplete lineage sorting, the average hidden-paralog rejection rate across all methods fell from 0.741 to 0.698. These results demonstrate how complete evolutionary ground truth enables direct benchmarking of a specific and otherwise difficult-to-measure source of orthology inference error.

**Figure 5:**
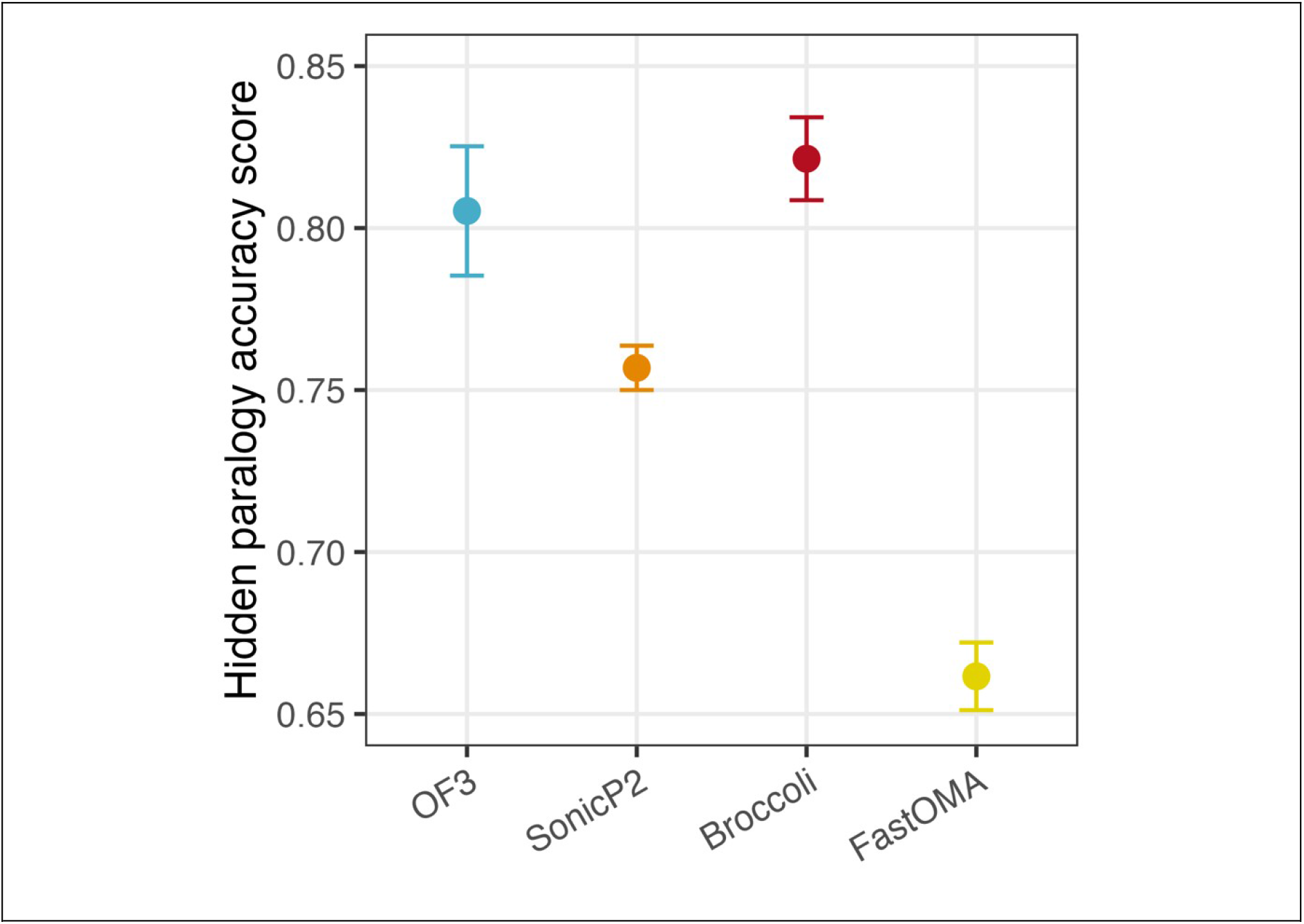
Hidden paralogy rejection rates differ between orthology inference methods. Hidden paralogy rejection rate for each tool on the QfO-trained simulations. A hidden paralog pair is a pair of genes from different species that are true paralogs, arising through differential loss following a duplication, but that share a simulated orthogroup and so appear orthologous. The hidden paralogy rejection rate is one minus the number of hidden paralog pairs a tool calls as orthologs, divided by the number of hidden paralog pairs it places within the same inferred orthogroup.

### Benchmarking is robust to the method used for calibration

Because OrthoSimulator is calibrated using empirically inferred orthogroups, an important potential source of bias is that an orthology method might perform better on simulations calibrated from its own output. We therefore tested explicitly for a self-calibration advantage. For each method, we compared its performance on the simulation calibrated from its own empirical orthogroups with its mean performance on simulations calibrated from the other three methods. Because the four simulated datasets differed slightly in overall difficulty, scores were first adjusted for these dataset-level differences. We then calculated a self-calibration delta, the difference between these two adjusted scores, for both orthogroup and ortholog F1, where positive values indicate better performance on a method’s own-calibrated simulation and values close to zero indicate little or no self-calibration advantage (Supplementary Figure 9).

Self-calibration effects were small for all four methods. For the QfO simulations, the largest positive effect was observed for FastOMA, with deltas of +0.024 for orthogroup F1 and +0.016 for ortholog F1. OrthoFinder showed no self-calibration advantage, with corresponding deltas of −0.007 and −0.001. The same overall pattern of small effects was observed for the OrthoBench simulations (Supplementary Figure 9). Thus, benchmark performance was not strongly dependent on which orthology method provided the empirical orthogroups used to calibrate the simulations.

### OrthoBench-calibrated simulations recapitulate empirical benchmark results

As an independent validation of the benchmarking framework, we tested whether simulations calibrated from the results of the orthology inference tools on the OrthoBench species reproduced the relative performance differences reported by the empirical OrthoBench benchmark. The published OrthoBench benchmark contains 70 expert-curated reference orthogroups, whereas each OrthoSimulator dataset contains 10,000 simulated orthogroups. Moreover, the 70 OrthoBench orthogroups are not a random sample, as some were deliberately selected to represent known challenges for orthology inference (Trachana et al. 2011). We therefore tested for broad agreement in orthology inference tool ranking rather than expecting an exact correspondence.

For each of the seven benchmark measures, we repeatedly sampled 70 reference orthogroups from the simulated dataset and compared the direction of each pairwise difference between methods with that reported by OrthoBench. In other words, we asked whether the method that scored higher in OrthoBench also scored higher in the simulation. Across the 36 pairwise comparisons for which the published benchmark showed a difference between methods, the median agreement was 87% across 1,000 resamples, and 25 of the 36 comparisons agreed with the published direction in most resamples (Supplementary Table 2). Thus, despite being generated without access to the curated OrthoBench reference orthogroups themselves, the simulated benchmark broadly recapitulated the relative performance differences observed in the empirical benchmark.

### OrthoSimulator reproduces empirical properties not used for calibration

Several properties examined above are either used directly during parameter estimation or contribute to the genetic algorithm fitness function. We therefore performed an additional held-out validation using three properties that were not used as simulation inputs or optimisation targets: sequence length, copy-number entropy across species, and pairwise sequence similarity. These properties instead emerge from the combined effects of the gene-tree, duplication and loss, and sequence-evolution models.

Across simulations calibrated from all four orthology methods and both empirical datasets, QfO and OrthoBench, the distributions of sequence length and copy-number entropy fell within the range of variation observed between empirical orthology methods. Pairwise sequence similarity showed a modest systematic difference, with simulated sequences tending to be more similar to one another than empirical sequences (Supplement S2). Thus, OrthoSimulator reproduces several empirical properties that were not explicitly used during calibration, while also identifying sequence similarity as one aspect where the current simulations remain less representative of empirical data.

## Discussion

Accurate benchmarking of orthology inference is essential for identifying limitations of existing methods and guiding future methodological development. The empirical resources developed by the Quest for Orthologs community provide a leading example of collaborative benchmarking grounded in real biological data and combining multiple complementary tests of orthology inference (Altenhoff et al. 2016; Linard et al. 2021; Nevers et al. 2022). Here, we present OrthoSimulator, a complementary simulation framework that generates orthogroup datasets calibrated from empirical data while retaining complete evolutionary ground truth. This provides scalable, realistic, and adaptable benchmark datasets where the underlying evolutionary relationships between orthologs and orthogroups can be examined.

OrthoSimulator enables direct evaluation of missing genes, orthogroup fission and fusion, pairwise ortholog inference and hidden paralogy. This extends benchmarking beyond orthogroup recovery to errors in individual orthologous relationships, evaluated against complete known ground truth. These properties can only be assessed partially in empirical datasets but remain important targets for continuing method development (Majidian, Hadziahmetovic, et al. 2025). Because OrthoSimulator also records the complete evolutionary history of every simulated orthogroup, individual inference errors can be traced back to the properties and evolutionary events associated with them. This creates opportunities to move beyond summary performance measures and investigate why particular classes of orthogroups are difficult to infer correctly.

To demonstrate the framework, we calibrated OrthoSimulator using orthogroups inferred from the species sets underlying two established benchmarking resources, QfO (Altenhoff et al. 2016) and OrthoBench (Trachana et al. 2011; Emms and Kelly 2020), and used the resulting simulations to evaluate four widely used orthology inference methods. The simulations reproduced many important properties of the empirical orthogroups, including orthogroup size, species representation, duplication number, and evolutionary rate. Benchmarking against the resulting ground truth revealed distinct error profiles among the four methods. Of the four methods tested, OrthoFinder was the most accurate overall for both orthogroup and pairwise ortholog inference, while other methods performed strongly for particular measures, including hidden-paralog rejection.

A potential concern with empirical calibration is that simulations could inherit biases from the orthology method used to generate the input orthogroups. We therefore calibrated independent simulations from four different methods and benchmarked every method across datasets derived from both its own and other methods’ outputs. Performance was consistent across independently calibrated simulations, with little evidence that methods gained an advantage when tested on simulations calibrated from their own output. Simulations calibrated from the OrthoBench species set also broadly recapitulated the relative performance differences observed in the independently curated OrthoBench reference orthogroups. In addition, held-out validation showed that the simulations reproduced empirical distributions of sequence length and copy-number entropy despite these properties not being used during calibration. Together, these analyses indicate that OrthoSimulator can reproduce the properties of empirical orthogroups without the benchmarking results being determined by the particular inference method used to calibrate the simulation.

These simulations may also have applications beyond benchmarking orthology inference itself. Complete evolutionary ground truth can be used to benchmark reconstruction of gene gain, loss, and duplication across species trees (Belcher and Kelly 2026), to evaluate species-tree methods that explicitly accommodate gene duplication and loss, such as ASTRAL-Pro (Zhang and Mirarab 2022), and to test functional annotation approaches that depend on orthology inference (Huerta-Cepas et al. 2017; Kirilenko et al. 2023). Efforts to reconstruct the gene repertoires of deep ancestral nodes, such as the last eukaryotic common ancestor (Richards et al. 2024), similarly depend on accurate reconstruction of gene families and could benefit from realistic simulated datasets in which the underlying evolutionary history is known.

Like any simulation-based approach, OrthoSimulator necessarily makes simplifying modelling assumptions. One important choice concerns incomplete lineage sorting. ILS can generate discordance between gene trees and species trees and may affect orthology inference in some evolutionary settings (Pamilo and Nei 1988; Degnan and Rosenberg 2006; Degnan and Rosenberg 2009; Majidian, Hadziahmetovic, et al. 2025). However, the magnitude of ILS appropriate for a particular empirical dataset cannot generally be estimated from orthogroup assignments alone. We therefore used a no-ILS baseline for the principal analyses rather than imposing an arbitrary level of coalescent discordance. Simulation frameworks such as SimPhy explicitly model ILS using a three-tree approach (Rasmussen and Kellis 2012; Mallo et al. 2016), but require additional demographic and evolutionary parameters that may not be directly estimable from the empirical orthogroup data used for calibration. OrthoSimulator instead provides an optional multispecies-coalescent extension in which ILS can be introduced using a single population-scaled parameter. This allows users to examine the consequences of ILS explicitly when appropriate without making it a fixed assumption of the empirically calibrated baseline. In our demonstration, introducing ILS altered absolute performance on several measures, particularly orthogroup fission and recall, but retained the broad differences between methods observed in the baseline benchmark.

Other aspects of sequence simulation also remain imperfect. Held-out validation showed that simulated sequences were systematically more similar to one another than empirical sequences, identifying sequence evolution as one area in which future versions could further improve realism. Gap simulation presents a related challenge. Previous work has shown that empirical and simulated alignments can be distinguished using gap information alone even under sophisticated indel models, and that reproducing empirical gap profiles can substantially improve realism (Trost et al. 2024). OrthoSimulator partly addresses this by using empirical gap information when constructing simulated alignments, but further improvements in indel and gap simulation are likely to be valuable. The modular structure of OrthoSimulator should allow improved models of sequence evolution, indels, and other evolutionary processes to be incorporated as they become available.

The Quest for Orthologs Consortium has highlighted disagreement between orthology inference methods as an important outstanding challenge (Majidian, Hadziahmetovic, et al. 2025). OrthoSimulator adds a complementary axis to existing orthology benchmarking by providing empirically calibrated simulations with complete evolutionary ground truth. While empirical benchmarks remain fundamental for evaluating methods against real biological data, simulated benchmarks allow controlled tests of the evolutionary conditions under which inference methods succeed or fail. By revealing exactly which orthogroups and orthologous relationships are recovered correctly, split, fused, missed, or confused by hidden paralogy, OrthoSimulator allows these differences to be measured and investigated directly. We release the simulated datasets, proteomes, and associated ground truth so that additional methods can be evaluated against the same reference data (figshare/zenodo link). Used alongside empirical resources such as QfO and OrthoBench, OrthoSimulator provides a framework for diagnosing inference errors, developing and testing new methods, and extending community benchmarking to questions for which empirical ground truth is unavailable.

## Methods

### Empirical datasets

We use two empirical datasets that underpin widely used orthology inference tools – Quest for Orthologs (QfO (Nevers et al. 2022)) and OrthoBench (Trachana et al. 2011; Emms and Kelly 2020). For QfO we downloaded the 2022 reference proteomes from the European Bioinformatics Institute (EBI) ftp site, using the 2022_02 with updated UP000000437 version. This dataset consists of 78 species (n=23 bacteria, n=7 archaea, n=48 eukaryotes). We used the same species tree as was used in the OrthoFinder v3 paper. For OrthoBench we obtained the 12 proteomes and the species tree from the supplementary material of Emms et al. 2020 (Emms and Kelly 2020).

### Empirical orthogroup inference

For each empirical dataset, we inferred orthogroups from the reference proteomes with four orthology inference methods. We ran OrthoFinder (Emms et al. 2026) version 3.1.3 (orthofinder -f <proteomes>). We ran FastOMA (Majidian, Nevers, et al. 2025) version 0.4.0 through its Nextflow pipeline using the conda execution profile, forcing pairwise ortholog generation and using the LUCA omamer DB (nextflow run dessimozlab/FastOMA -r v0.4.0 -profile conda --input_folder <proteomes> --output_folder <output> --omamer_db LUCA.h5 --force_pairwise_ortholog_generation true). We ran SonicParanoid2 (Cosentino et al. 2024) in its default mode (sonicparanoid -i <proteomes> -o <output> -m default). We ran Broccoli (Derelle et al. 2020) with default settings (python broccoli.py -dir <proteomes>). Each of the four resulting sets of empirical orthogroups was used, in turn, to train an independent OrthoSimulator simulation for that dataset (see ‘Cross-validation design’ below).

For each simulated dataset, the distributions of key orthogroup properties were compared with those of the corresponding empirical orthogroups. The SonicParanoid2-calibrated QfO dataset is shown in Figure 2, with the remaining comparisons shown in Supplementary Figures 1–7. Differences between distributions were quantified using the 1-Wasserstein distance, where a value of zero indicates identical distributions. Because the absolute value of this distance depends on the property being measured, the simulation-to-empirical distance was compared with the distances observed between the four empirical orthology methods analysing the same genomes. We express this relative to the largest difference between any two empirical methods. Thus, a value of 100% would mean that the simulated and empirical distributions differ by as much as the two most divergent empirical methods, whereas values below 100% indicate closer agreement.

### Cross-validation design

For each empirical dataset (QfO and OrthoBench), we used OrthoSimulator to train four independent simulations, one from the empirical orthogroups inferred by each of the four orthology inference methods (OrthoFinder, FastOMA, SonicParanoid2, Broccoli). Each trained simulation comprised 10,000 simulated orthogroups distributed across the dataset’s proteomes, giving eight simulated datasets in total (four trained on QfO, four on OrthoBench). We then benchmarked each of the four orthology inference methods against all eight simulated datasets, so that every tool was tested on two simulations trained from its own empirical orthogroups and six trained from the other three methods. This design separates tool performance from dataset and training-method effects, lets us test directly for any advantage a tool gains from being benchmarked on data trained from its own output, and lets us report precision and recall with error bars by averaging each tool’s performance across the simulations trained on all four methods.

### Empirical parameter estimation

To train a simulation on real data, we took the sequences underlying a set of empirical orthogroups and estimated sequence-evolution, branch-rate, indel and alignment-structure parameters directly from them. We used these per-orthogroup estimates to build an empirical distribution for each parameter, from which OrthoSimulator draws a value independently for every orthogroup it simulates.

#### Orthogroup sampling

We sampled 5,000 orthogroups at random from each empirical orthogroup set, restricting sampling to orthogroups containing between 4 and 499 genes. This range excludes orthogroups too small to give a meaningful tree, while keeping alignment and tree inference fast enough to run at scale.

#### Multiple sequence alignment and gene tree inference

We aligned each sampled orthogroup with MAFFT (Katoh and Standley 2013) L-INS-i (--maxiterate 1000 --localpair), removed poorly aligned columns with trimAl’s (Capella-Gutiérrez et al. 2009) ‘gappyout’ option, and inferred a maximum-likelihood gene tree from the trimmed alignment with IQ-TREE3 (Wong et al. 2025) under a fixed JTT+G4+I model in fast search mode.

#### Sequence-evolution parameters

We extracted the proportion of invariant sites and the gamma shape parameter from each orthogroup’s IQ-TREE log, discarding gamma-shape estimates above 10 as implausible outliers. For each parameter, we trimmed the remaining per-orthogroup values to the 10th to 90th percentile, took the mean of the trimmed set, and estimated its standard deviation with a robust median absolute deviation estimator (MAD × 1.4826). We converted the trimmed mean and standard deviation to log-space with the delta method, giving log-normal shape and scale parameters that OrthoSimulator draws from independently for every simulated orthogroup.

#### Branch-rate heterogeneity

To parameterise branch-length relaxation (see ‘Branch-length relaxation’ below), we estimated how far branch-specific rates depart from the expectation under a strict molecular clock. For each sampled gene tree, we forced a copy of the tree to be ultrametric and compared it branch-by-branch with the original tree. For each branch, we calculated the relative rate as the ratio of its observed length to the corresponding ultrametric length, then took the standard deviation of log(rate) - log(parent rate) across all branches in the tree as a per-tree sigma. We trimmed sigma to the 5th to 95th percentile across orthogroups, log-transformed it, and used the mean and standard deviation of the log-transformed values to define a log-normal distribution. During simulation, OrthoSimulator draws a sigma value from this distribution and squares it before passing it to the branch-relaxation model.

#### Empirical gap-position profiles

We built one empirical gap-position profile per sampled orthogroup from its untrimmed MAFFT alignment. We divided the alignment into 100 equal-width bins along its length and recorded, for each bin, the proportion of that alignment’s total gap characters it contained. During simulation, OrthoSimulator selects one of these profiles at random for each orthogroup and uses it to reposition simulated gap blocks (see ‘Gap-position shuffling’ below).

#### Species tree rescaling

We rescaled the input species tree so that its root-to-tip distance matched the median of the per-orthogroup median root-to-tip distances from the sampled empirical trees, then re-imposed the ultrametric constraint. We used this rescaled tree as the host tree for all subsequent gene-tree simulations.

### PFAM domain model extraction

A domain-aware simulation needs three things for a given proteome, 1) PFAM domain annotations, 2) a mapping from each protein to its domains, and 3) a fitted substitution model for each domain. We provide these precomputed for twelve model species (see ‘The OrthoSimulator workflow’ in Results) and provide a module that builds them for any other proteome.

To build these models, we scanned each proteome with pfam_scan.pl (Finn et al. 2010) against the Pfam-A HMM database, keeping only hits of type Domain. For each unique PFAM domain identified, we downloaded its seed alignment from InterPro (Blum et al. 2025) in Stockholm format, converted it to FASTA, and fitted a substitution model with IQ-TREE3, restricting model selection to LG+I and JTT+G with up to four rate or mixture classes (-mset LG+I,JTT+G --cmax 4 --fast). From each resulting IQTree log we parsed the best-fit model, proportion of invariant sites, gamma shape parameter and any free-rate weights, and assembled these into an AliSim-compatible model string (e.g. JTT+I{0.12}+G4{0.87}).

During simulation, OrthoSimulator assigns a domain region the model drawn at random from this species-wide table of fitted domain models (see ‘Domain-aware sequence simulation’ below).

### Orthogroup simulation

We simulated each orthogroup independently, with no protein used to seed more than one orthogroup.

#### Gene tree simulation

We simulated each gene tree inside the rescaled species tree with SaGePhy’s (Kundu and Bansal 2019) GuestTreeGen, under a birth-death process allowing duplication, loss and horizontal transfer (java -jar sagephy.jar GuestTreeGen -a 250 -gb -gbc <gbc> -vp XTAGX -p <leaf_sampling_probability> -rt <replacement_prob> <species tree> <duplication rate> <loss rate> <transfer rate> <output>). For each orthogroup, we drew the duplication and loss independently from uniform distributions bounded by zero and their respective trained maxima. The orthogroup-birth-bias parameter gbc controls where on the species tree new gene families tend to arise, with lower values pushing new families towards the tips, mimicking younger genes.

#### Incomplete lineage sorting (optional)

SaGePhy merges gene lineages exactly at each speciation, so it cannot produce incomplete lineage sorting (ILS) by itself. To let users test the effect of ILS, we added an optional step that rearranges each gene tree after SaGePhy has simulated it. The rearrangement follows the multispecies coalescent (Rannala and Yang 2003; Degnan and Salter 2005). A single parameter, theta (--ils-theta), sets the amount of ILS, and lower values give more discordance between gene trees and the species tree. Duplications keep their simulated dates, and copies descended from a duplication coalesce no later than that duplication (the bounded coalescent; Rasmussen and Kellis 2012). We use the rearranged tree to simulate sequences. We still read true orthologs from the original SaGePhy tree, so the ground truth does not change. ILS is off unless the user sets theta. All baseline simulations in this paper use no ILS, and the ILS benchmark (Supplementary Figure 10) uses theta = 10.

#### Branch-length relaxation

Because real gene trees rarely evolve under a strict clock, we relaxed the branch lengths of every simulated gene tree with SaGePhy’s BranchRelaxer under the ACRY07 autocorrelated-rate model (java -jar sagephy.jar BranchRelaxer <pruned tree> ACRY07 <start rate> <sigma> --keep-interior-names -o <output>). We fixed the starting rate at 1 and drew sigma independently per orthogroup from the log-normal distribution estimated above, squaring it before passing it to BranchRelaxer.

#### Starting sequence selection and age-dependent shuffling

We seeded each orthogroup with one real protein sequence drawn without replacement from the chosen seed species’ proteome. If the orthogroup’s origin node doesn’t cover >=90% of all species, we shuffled the seed sequence in 20-residue blocks.

#### Domain-aware sequence simulation

Before simulating sequence evolution, we built a partition file dividing the seed sequence into domain and non-domain regions, merging any overlapping PFAM hits into single intervals. We assigned non-domain regions the background model JTT+G4{alpha}+I{p_inv}. We assigned each domain region a model drawn at random from the seed species’ PFAM model table, adding the C20 empirical profile mixture whenever the underlying model was LG or JTT.

We simulated sequence evolution along the relaxed, relabelled gene tree with IQ-TREE3’s AliSim (Ly-Trong et al. 2022) module, supplying the partition file, tree and seed sequence (iqtree3 --quiet --alisim <output> --indel <insertion rate>,<deletion rate> --indel-size POW{1.7/50},POW{2.2/40} -p <partition file> -t <tree> --out-format fasta --root-seq <seed>, <name>). We drew the insertion and deletion rates independently per orthogroup from uniform distributions bounded between 0-0.04 and drew indel lengths using an asymmetric power law.

#### Leaf relabelling

After simulation, we relabelled every leaf as <species>_<orthogroup>_<copy number> (for example, Homo_sapiens_12_2 is the second human copy in simulated orthogroup 12. This mapping lets downstream steps assign genes to species and recover true ortholog pairs.

#### Gap-position shuffling

AliSim distributes indels fairly uniformly along an alignment, but real protein alignments tend to concentrate gaps in structurally flexible regions. To correct this, we identified contiguous gapped column blocks in each simulated alignment and ranked them from least to most gap-rich. We then picked one empirical gap-position profile at random, ranked its 100 positional bins the same way, and reassigned simulated blocks so that the gappiest simulated blocks moved towards the positions ranked as gap-rich in the chosen empirical profile. Columns without any gaps kept their original order throughout, so alignment length and total gap content were unchanged.

#### Quality control

We checked every simulated alignment for sequences made up entirely of gap characters, which can arise for very short or highly diverged simulated genes. Where this occurred, we reran the simulation from scratch using the same seed sequence, repeating until we obtained a gap-free alignment.

#### Recording the true evolutionary history

For every accepted orthogroup, we read SaGePhy’s guest-to-host mapping file and recorded the true number of duplication and loss events on the gene tree. We identified true ortholog pairs by traversing the gene tree and, at every node representing a speciation event, recording all pairs of genes descending from its two child lineages; we classified gene pairs separated by a duplication node as paralogs and excluded them from the ortholog set.

### Genetic algorithm parameter optimisation

To estimate the rates of gene duplication, loss and gbc (The orthogroup-birth-bias parameter) a genetic algorithm was employed. As these parameters are complementary multiple possible solutions can exist that satisfy the empirical data. For example, high duplication with high loss may mimic low loss and low duplication scenarios. Normal traversal programme methods such as binary search methods may exclude good solutions. The genetic algorithm was implemented using PyGad, a python library for building genetic algorithms, requiring an empirical output and empirical parameter estimation file (mentioned previously). We provide flexible parameter optimisation for users to decide on key parameters for the GA. These are the number of generations, parent mating, solutions per population, parent selection type, crossover probability, mutation type, mutation percentage, sample size and number of Orthogroups, further information about each PyGad specific parameter can be found on the PyGad GitHub.

During each generation each member of the population is tested against the empirical distributions using a fitness function. The parameters for gene duplication, loss and gbc from each individual are simulated for some number of orthogroups with those parameters derived directly from the empirical data. Then each simulation is compared to feature distributions from the empirical data; the number of genes per Orthogroup, number of species per Orthogroup, and the duplication counts. This involves deriving the Kullback-Leibler (KL) divergence between identical bins (derived using Freedman Diaconis Estimator), a large KL value indicates that the distributions are highly divergent. This distance parameter is then converted to a fitness value by taking the reciprocal (1/1+KL) with high divergence signifying low fitness. The genetic algorithm then runs for the user specified number of iterations. Following this it then reports the solution for duplication, loss and gbc with the peak fitness value and the final iteration solution, producing config files containing these values in addition to the previously calculated parameters allowing users to immediately run these simulations. The fitted values for each calibration are given in Supplementary Table 3.

### Quantitative assessment of simulation realism

To assess how closely each simulated dataset reproduced its training data, we calculated the 1-Wasserstein distance between the simulated and empirical distributions for eight curated orthogroup properties: number of genes, number of species, number of duplications, gamma shape parameter, proportion of invariant sites, median root-to-tip distance, treeness, and number of gap regions per alignment. Each distance was approximated from a grid of 2,000 matched quantiles per distribution. To establish a reference scale for this distance, we also inferred orthogroups from the same reference proteomes with all four orthology methods used for training (OrthoFinder, FastOMA, Broccoli, SonicParanoid2; see ‘Empirical orthogroup inference’ above), and calculated the same Wasserstein distance between every pairwise combination of these four empirical outputs, for each of the eight properties. We report the simulation-training distance as a ratio to the mean and to the maximum of these six pairwise empirical distances, rather than applying a fixed pass or fail threshold. With only six pairwise comparisons available, a hard cutoff, for example requiring the simulation to be no more divergent than the single most divergent empirical pair, is highly sensitive to which pair happens to be the most divergent. A ratio at or below 1 indicates that the simulation diverges from its training data no more than two independently run real orthology methods already diverge from each other on the same genomes; ratios above 1 are reported directly rather than folded into a threshold. This analysis was carried out for each of the four training tools on the QfO and OrthoBench datasets (Figure 2, Supplementary Figures 1–7, Supplementary Table 1).

### Benchmarking orthogroup and ortholog inference

We assessed the accuracy of each orthology inference method at three complementary levels: orthogroup recovery, ortholog-pair recovery, and hidden paralogy.

#### Orthogroup-level benchmarking

We compared inferred orthogroups against the complete set of simulated reference orthogroups, first removing any predicted orthogroup containing only a single gene. For each reference orthogroup, we calculated the fraction of its genes recovered by each overlapping predicted orthogroup (recall) and the fraction of each predicted orthogroup’s genes that belonged to the reference orthogroup (precision), combined these across all overlapping predicted orthogroups weighted by overlap size, then averaged recall and precision across reference orthogroups weighted by reference-orthogroup size. We calculated their harmonic mean as F1. We defined fusion as the percentage of reference orthogroups whose genes appeared, alongside genes from at least one other reference orthogroup, in the same predicted orthogroup, and fission as the percentage of reference orthogroups whose genes were split across more than one predicted orthogroup. We defined missing genes as the percentage of reference-orthogroup genes absent from every predicted orthogroup, and missing reference orthogroups as the percentage of reference orthogroups with no genes in any predicted orthogroup. For entropy, we built, for each reference orthogroup, a distribution over the predicted orthogroups its genes were assigned to, calculated its Shannon entropy in bits, normalised this by log2(orthogroup size), and averaged across reference orthogroups weighted by size. To produce a single composite score of orthogroup-inference accuracy for each tool, we converted each of these seven measures to a z-score against the across-tool mean and summed them.

#### Ortholog-pair benchmarking

We compared each tool’s inferred ortholog pairs against the complete set of true simulated pairs, excluding same-species pairs from the true set, and defined true positives as correctly recovered true pairs, false positives as inferred pairs that are not true orthologs, and false negatives as true pairs the tool did not recover. We calculated precision, recall and their harmonic mean (F1) for each tool.

#### Hidden paralogy benchmarking

We defined a tool’s hidden-paralogy rejection rate as one minus the number of false-friend pairs a tool calls as orthologs divided by the number of false-friend pairs it places within the same inferred orthogroup (the pairs it had the opportunity to call orthologs).

## Acknowledgements

L.B., J.H. and S.K. were funded by the Wellcome Trust under grant agreement number 226598/Z/22/Z.

## Data availability

OrthoSimulator is open-source software released under the GNU General Public License v3.0 and is available at https://github.com/lauriebelch/OrthoSimulator. The eight simulated datasets, the parameter estimates they were calibrated from, all benchmark results, and the scripts that regenerate figures and supplementary tables are available on figshare (https://doi.org/10.6084/m9.figshare.34988026).

## Author contributions

L.B. and S.K. conceived the study. L.B. led the study and designed the overall OrthoSimulator framework and benchmarking strategy. J.H. contributed to study design and methodological development throughout the project, co-developed the OrthoSimulator software, and designed and implemented the genetic algorithm module. L.B. performed the empirical orthogroup inference, simulations, benchmarking, and analyses, and supervised the project and software development. L.B. wrote the original manuscript, with contributions from J.H. and S.K. All authors reviewed and edited the manuscript.

