## Supplementary Info for "OrthoSimulator: Empirically trained simulations for ground-truth benchmarking of orthology inference"

### OrthoSimulator: Supplementary Material

This supplement extends the main text with seven additional realism comparisons covering the calibration methods and dataset not shown in Figure 2 (Supplementary Figures 1 to 7, Supplementary Table 1), orthogroup-level, ortholog-pair, and hidden-paralogy benchmarking results for the OrthoBench dataset extending Figures 3 to 5 (Supplement S1), a schematic illustration of hidden paralogy (Supplementary Figure 8), the self-calibration analysis referenced in the main text (Supplementary Figure 9), validation of the OrthoBench-calibrated simulations against the independently published OrthoBench benchmark (Supplementary Table 2), a held-out validation panel using three measures that play no role in simulation calibration or parameter optimisation (Supplement S2), a supplementary benchmark run under simulated incomplete lineage sorting (Supplementary Figure 10), and the genetic-algorithm-optimised simulation parameters underlying all main-text results (Supplementary Table 3).

### Realism comparisons across calibration methods and datasets

Figure 2 in the main text compares the SonicParanoid2-calibrated QfO simulation against its own calibration data across eight orthogroup and gene-tree properties. Supplementary Figures 1 to 7 below show the same eight-panel comparison for each of the remaining calibration method and dataset combinations, OrthoFinder, FastOMA, and Broccoli calibrated from QfO, and all four tools calibrated from OrthoBench. Supplementary Table 1 reports the accompanying quantitative realism check, for the QfO dataset and then the OrthoBench dataset. For each measure and calibration method, we computed the Wasserstein distance between the simulated and empirical calibration distributions, and expressed it as a ratio to the mean and maximum of the six pairwise Wasserstein distances among the four empirical tools' own outputs on the same measure. A ratio near 1 means the simulation is about as far from its calibration data as two real tools already are from each other on that measure.

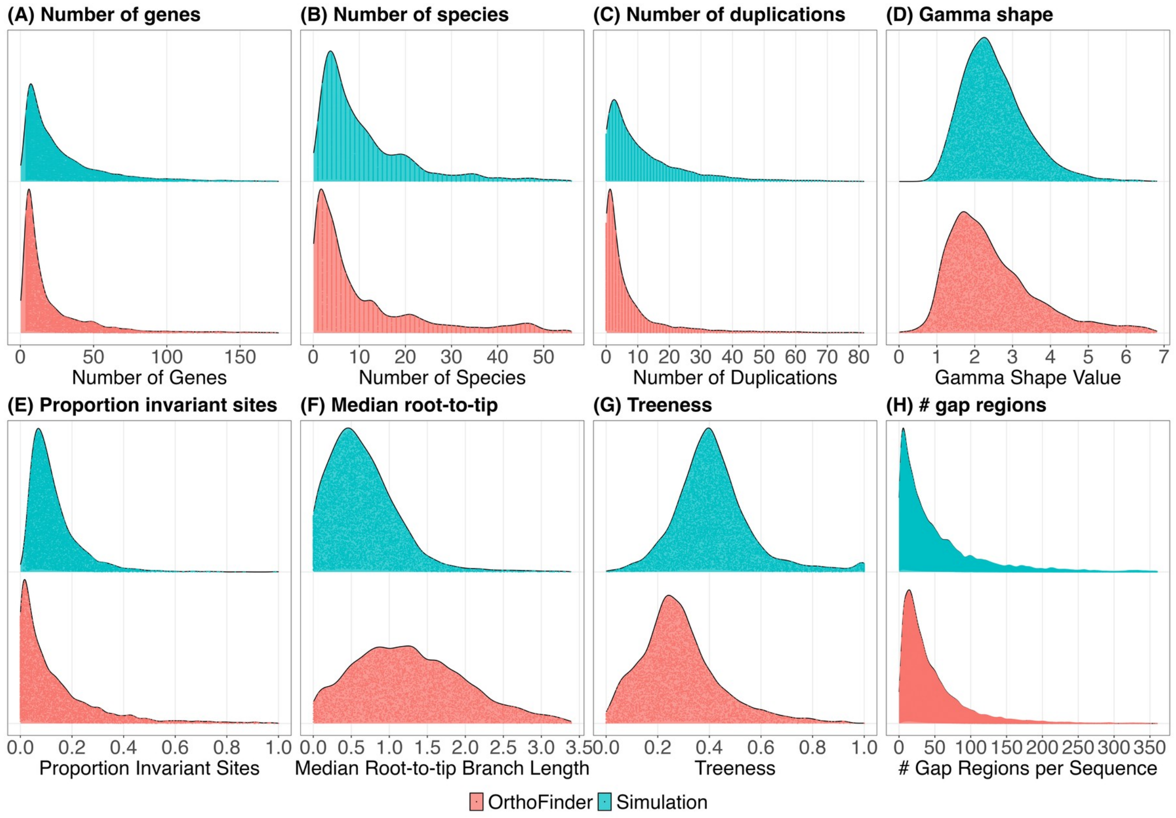


**Supplementary Figure 1**. Realism comparison for QfO dataset calibrated from OrthoFinder. Distributions of eight orthogroup properties for the OrthoFinder-calibrated QfO simulation (blue) compared against the empirical OrthoFinder orthogroups used to calibrate it (red). Data shown for (A) Number of genes per orthogroup. (B) Number of species represented per orthogroup. (C) Number of duplications per orthogroup. (D) Gamma shape parameter of the among-site rate-variation model. (E) Proportion of invariant sites per alignment. (F) Median root-to-tip distance per gene tree. (G) Treeness, the proportion of total branch length found on internal branches. (H) Number of gap regions per aligned sequence.

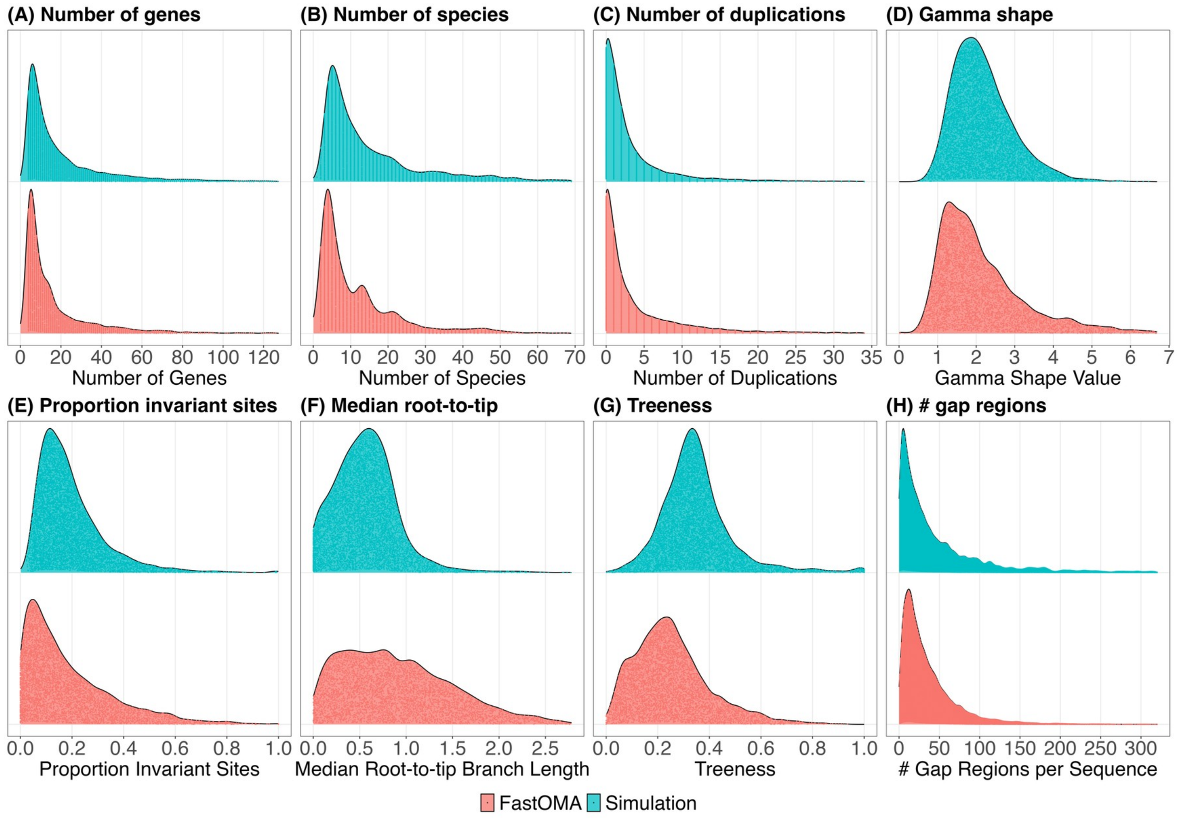


**Supplementary Figure 2**. Realism comparison for QfO dataset calibrated from FastOMA. Panels (A) to (H) same as Supplementary Figure 1.

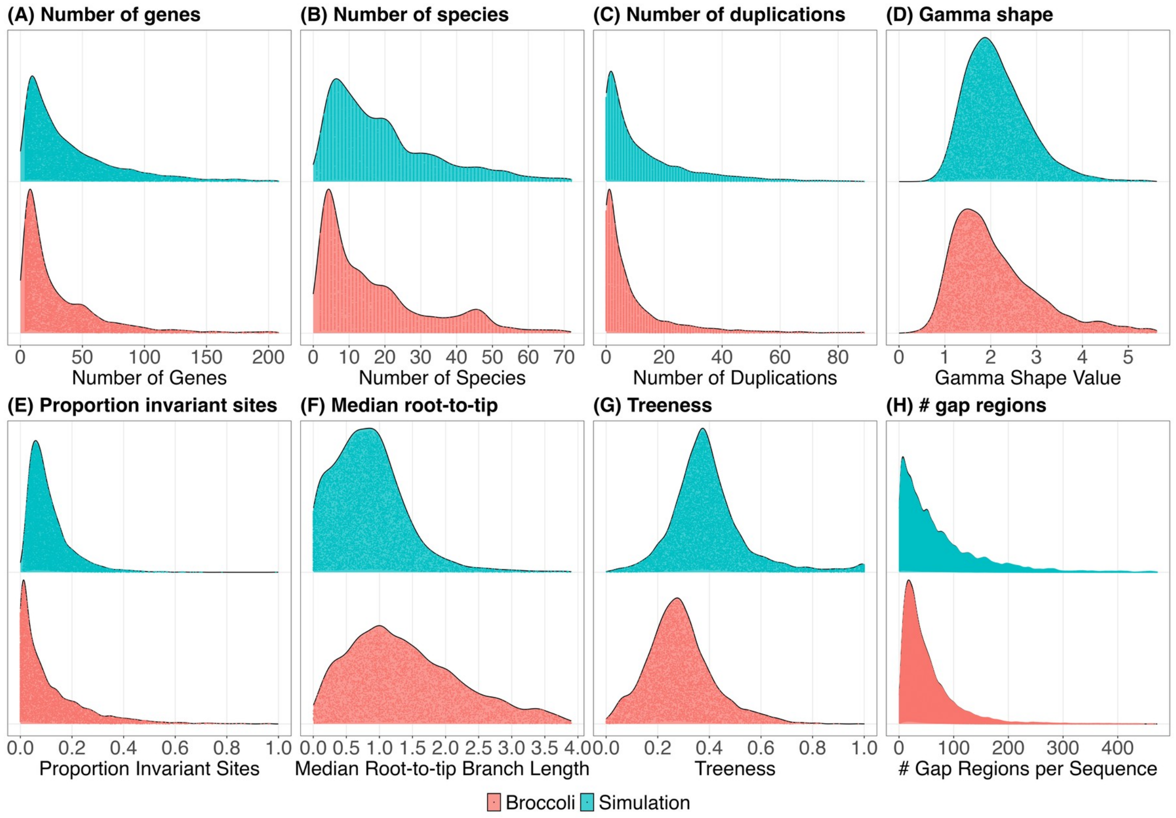


**Supplementary Figure 3**. Realism comparison for QfO dataset calibrated from Broccoli. Panels (A) to (H) same as Supplementary Figure 1.

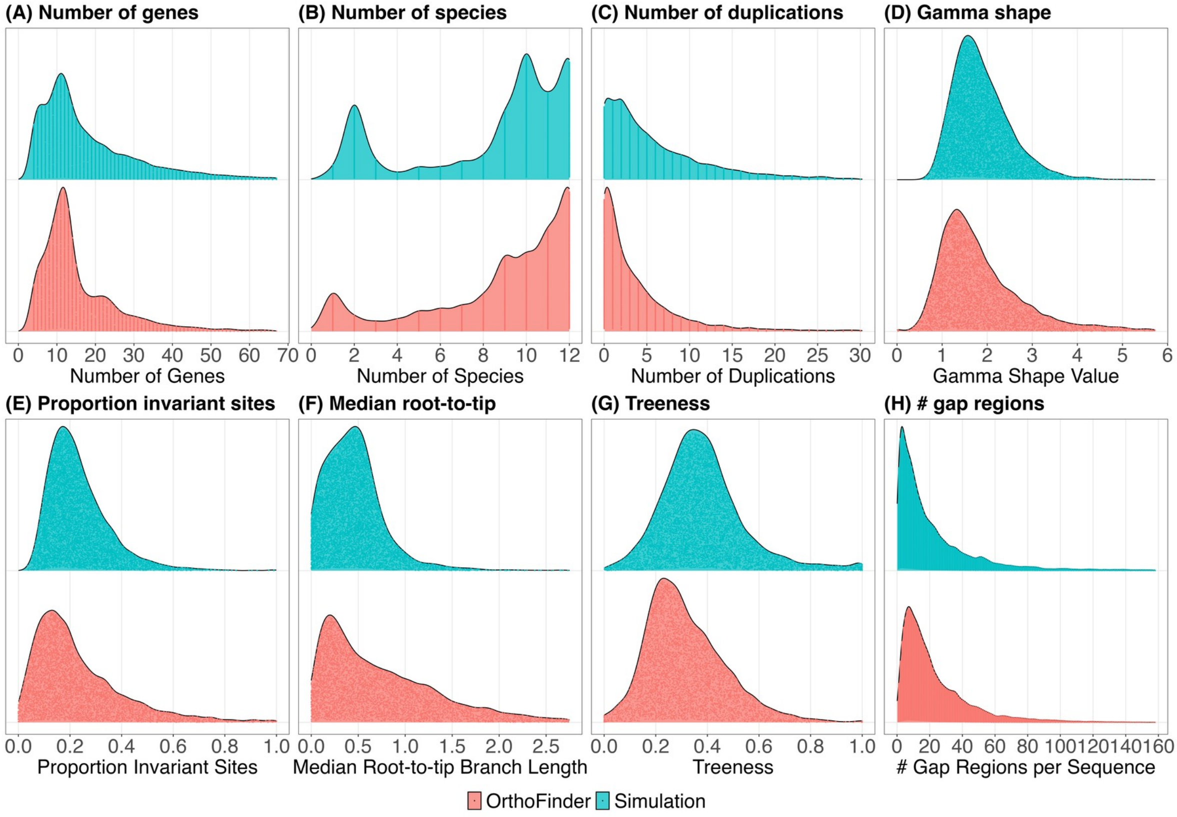


**Supplementary Figure 4**. Realism comparison for OrthoBench dataset calibrated from OrthoFinder. Panels (A) to (H) same as Supplementary Figure 1.

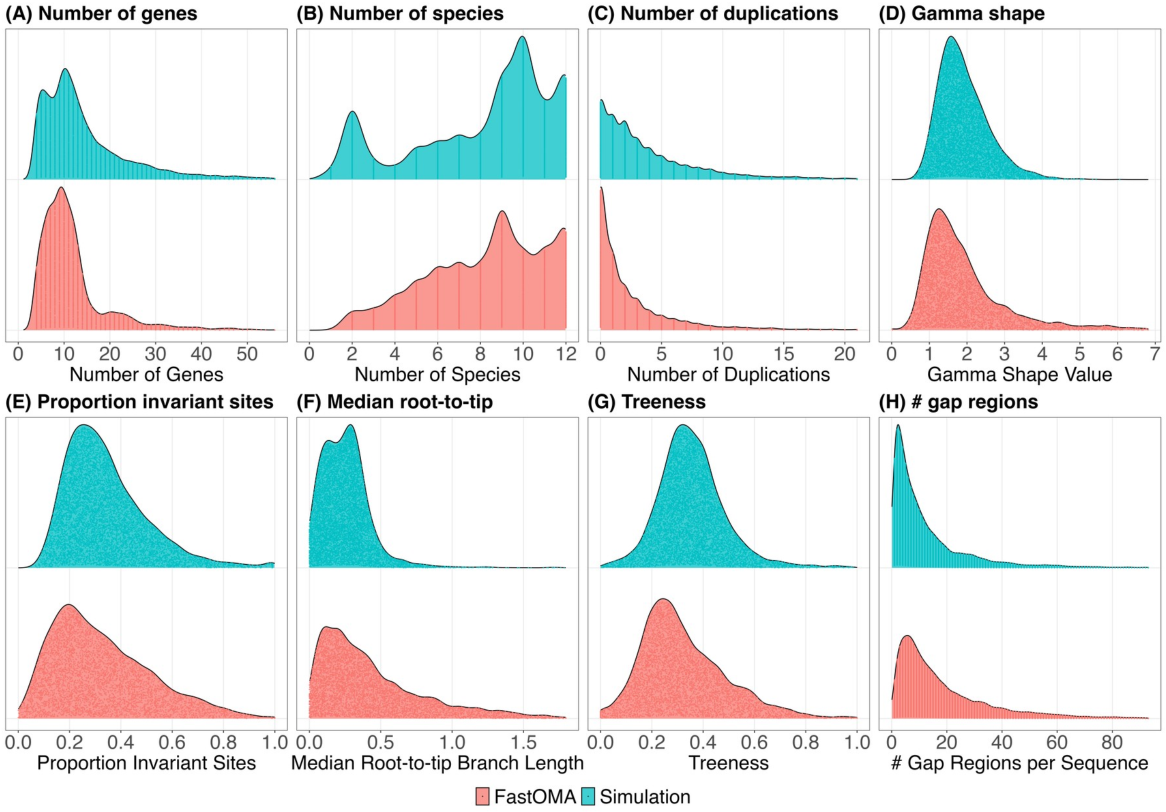


**Supplementary Figure 5**. Realism comparison for OrthoBench dataset calibrated from FastOMA. Panels (A) to (H) same as Supplementary Figure 1.

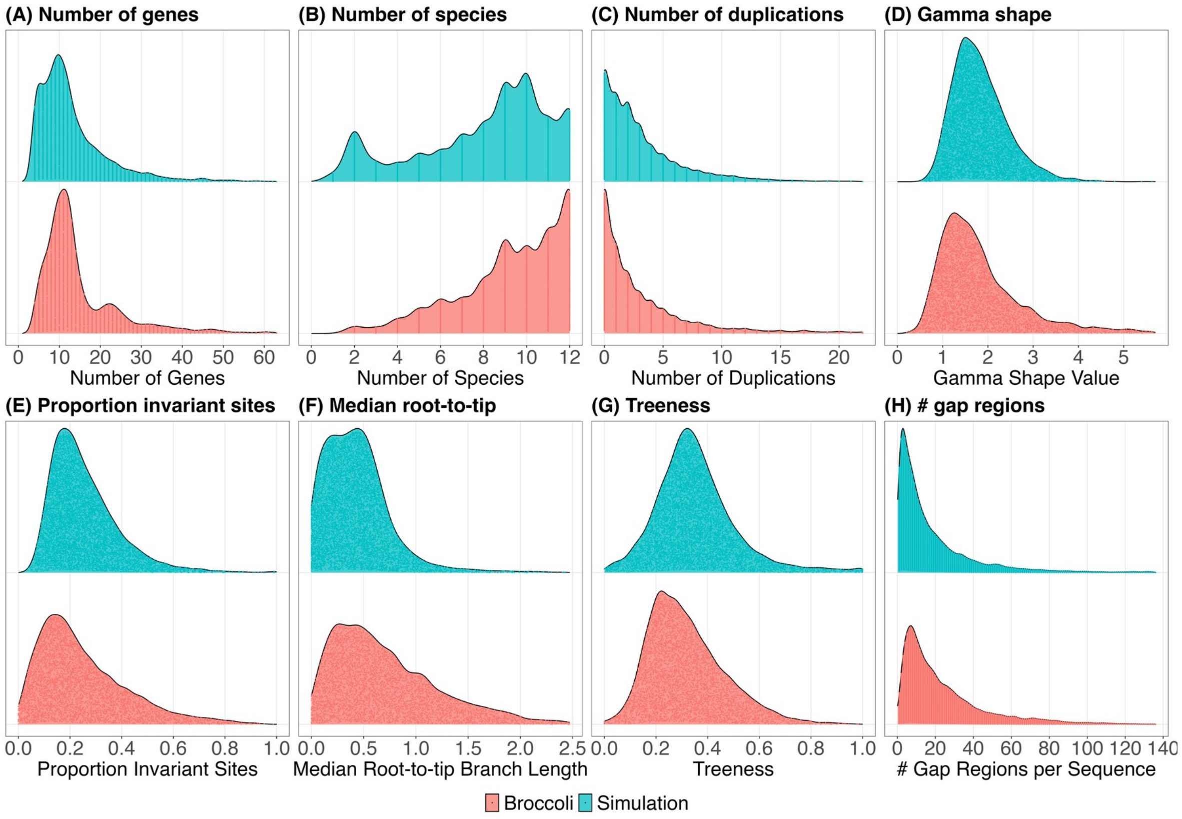


**Supplementary Figure 6**. Realism comparison for OrthoBench dataset calibrated from Broccoli. Panels (A) to (H) same as Supplementary Figure 1.

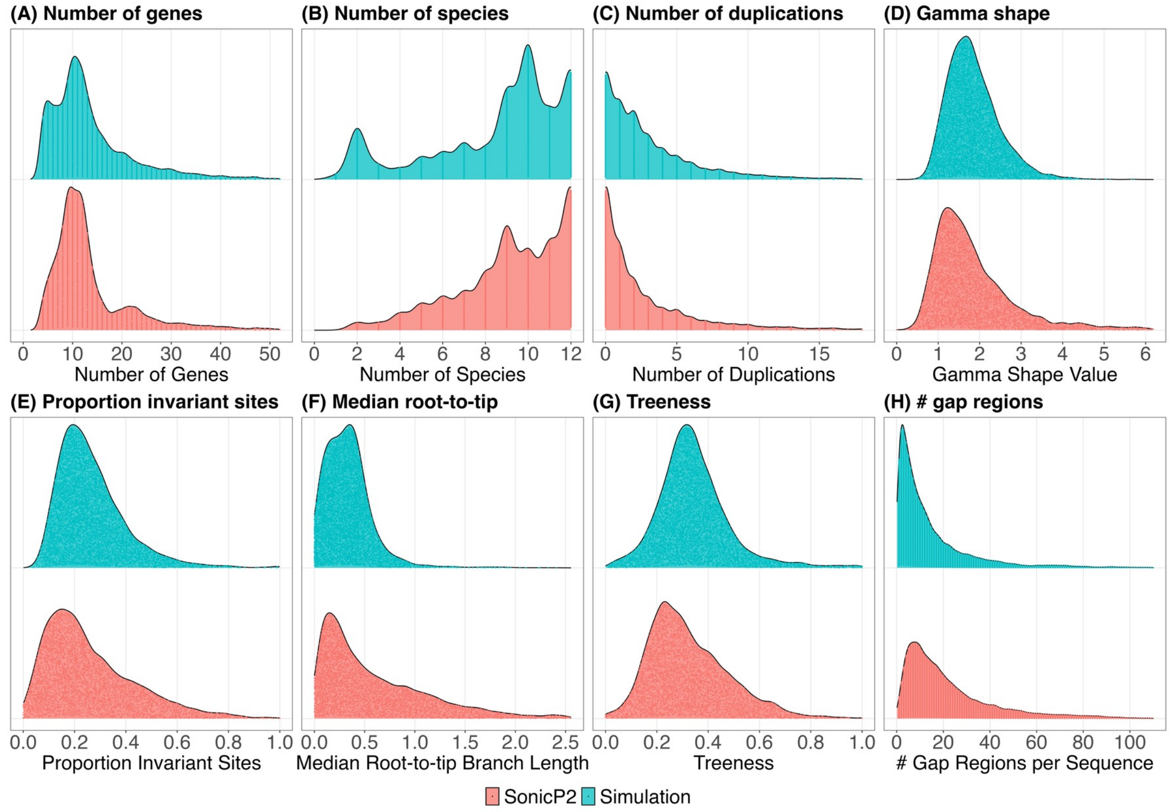


**Supplementary Figure 7**. Realism comparison for OrthoBench dataset calibrated from SonicParanoid2. Panels (A) to (H) same as Supplementary Figure 1.

**Supplementary Table 1**. Quantitative realism check for the QfO and OrthoBench datasets. The value W1 (sim, empirical) is the Wasserstein distance between the simulated and empirical calibration distributions for that measure. The two ratio columns express this relative to the mean and maximum of the six pairwise Wasserstein distances among the four empirical tools' outputs. This demonstrates how different the simulation is from the empirical distribution, compared to how different the empirical distributions from different tools are from each other.

**QfO dataset**

| **Calibrated from** | **Measure** | **W1 (sim, empirical)** | **Ratio to empirical mean** | **Ratio to empirical max** |
|---|---|---|---|---|
| OrthoFinder | num_genes | 7.46 | 0.84 | 0.44 |
| OrthoFinder | num_species | 2.16 | 0.54 | 0.36 |
| OrthoFinder | num_duplications | 6.77 | 1.75 | 1.03 |
| OrthoFinder | gamma_shape | 0.41 | 1.91 | 1.10 |
| OrthoFinder | proportion_invariant | 0.05 | 1.15 | 0.61 |
| OrthoFinder | median_rtt | 0.67 | 2.45 | 1.29 |
| OrthoFinder | treeness | 0.13 | 5.53 | 3.65 |
| OrthoFinder | num_gaps | 15.59 | 1.85 | 1.09 |
| FastOMA | num_genes | 3.40 | 0.38 | 0.20 |
| FastOMA | num_species | 4.10 | 1.03 | 0.69 |
| FastOMA | num_duplications | 1.85 | 0.48 | 0.28 |
| FastOMA | gamma_shape | 0.40 | 1.89 | 1.08 |
| FastOMA | proportion_invariant | 0.04 | 1.08 | 0.57 |
| FastOMA | median_rtt | 0.37 | 1.38 | 0.72 |
| FastOMA | treeness | 0.09 | 4.04 | 2.67 |
| FastOMA | num_gaps | 15.07 | 1.79 | 1.05 |
| Broccoli | num_genes | 8.79 | 0.99 | 0.52 |
| Broccoli | num_species | 2.84 | 0.71 | 0.48 |
| Broccoli | num_duplications | 5.84 | 1.51 | 0.89 |
| Broccoli | gamma_shape | 0.35 | 1.62 | 0.93 |
| Broccoli | proportion_invariant | 0.04 | 0.97 | 0.51 |
| Broccoli | median_rtt | 0.62 | 2.26 | 1.19 |
| Broccoli | treeness | 0.11 | 4.69 | 3.10 |
| Broccoli | num_gaps | 29.45 | 3.50 | 2.06 |
| SonicParanoid2 | num_genes | 5.95 | 0.67 | 0.35 |
| SonicParanoid2 | num_species | 5.24 | 1.31 | 0.88 |
| SonicParanoid2 | num_duplications | 3.89 | 1.01 | 0.59 |
| SonicParanoid2 | gamma_shape | 0.35 | 1.66 | 0.95 |
| SonicParanoid2 | proportion_invariant | 0.04 | 0.96 | 0.50 |

| Calibrated from | Measure | W1 (sim, empirical) | Ratio to empirical mean | Ratio to empirical max |
|---|---|---|---|---|
| SonicParanoid2 | median_rtt | 0.47 | 1.72 | 0.90 |
| SonicParanoid2 | treeness | 0.07 | 2.83 | 1.87 |
| SonicParanoid2 | num_gaps | 23.95 | 2.85 | 1.67 |

**OrthoBench dataset**

| Calibrated from | Measure | W1 (sim, empirical) | Ratio to empirical mean | Ratio to empirical max |
|---|---|---|---|---|
| OrthoFinder | num_genes | 2.88 | 1.05 | 0.64 |
| OrthoFinder | num_species | 0.50 | 0.69 | 0.47 |
| OrthoFinder | num_duplications | 3.70 | 2.66 | 1.53 |
| OrthoFinder | gamma_shape | 0.35 | 4.35 | 2.57 |
| OrthoFinder | proportion_invariant | 0.05 | 0.92 | 0.49 |
| OrthoFinder | median_rtt | 0.33 | 1.78 | 1.05 |
| OrthoFinder | treeness | 0.07 | 7.99 | 5.17 |
| OrthoFinder | num_gaps | 5.37 | 1.87 | 0.99 |
| FastOMA | num_genes | 2.67 | 0.97 | 0.60 |
| FastOMA | num_species | 0.50 | 0.69 | 0.47 |
| FastOMA | num_duplications | 2.70 | 1.94 | 1.12 |
| FastOMA | gamma_shape | 0.45 | 5.49 | 3.25 |
| FastOMA | proportion_invariant | 0.04 | 0.76 | 0.40 |
| FastOMA | median_rtt | 0.22 | 1.20 | 0.71 |
| FastOMA | treeness | 0.05 | 5.75 | 3.72 |
| FastOMA | num_gaps | 6.03 | 2.10 | 1.11 |
| Broccoli | num_genes | 3.68 | 1.34 | 0.82 |
| Broccoli | num_species | 1.31 | 1.82 | 1.23 |
| Broccoli | num_duplications | 2.20 | 1.58 | 0.91 |
| Broccoli | gamma_shape | 0.36 | 4.37 | 2.59 |
| Broccoli | proportion_invariant | 0.04 | 0.78 | 0.41 |
| Broccoli | median_rtt | 0.30 | 1.62 | 0.96 |
| Broccoli | treeness | 0.04 | 4.34 | 2.81 |
| Broccoli | num_gaps | 6.60 | 2.30 | 1.21 |
| SonicParanoid2 | num_genes | 1.69 | 0.61 | 0.38 |
| SonicParanoid2 | num_species | 0.64 | 0.89 | 0.60 |
| SonicParanoid2 | num_duplications | 2.15 | 1.55 | 0.89 |
| SonicParanoid2 | gamma_shape | 0.39 | 4.74 | 2.80 |

| Calibrated from | Measure | W1 (sim, empirical) | Ratio to empirical mean | Ratio to empirical max |
|---|---|---|---|---|
| SonicParanoid2 | proportion_invariant | 0.04 | 0.84 | 0.44 |
| SonicParanoid2 | median_rtt | 0.32 | 1.76 | 1.04 |
| SonicParanoid2 | treeness | 0.03 | 3.75 | 2.43 |
| SonicParanoid2 | num_gaps | 7.98 | 2.78 | 1.47 |

### Supplement S1: OrthoBench benchmarking results

Figures 3 to 5 in the main text report orthogroup-level, ortholog-pair, and hidden-paralogy benchmarking results for the four orthology inference methods on the QfO-calibrated simulations. This supplement reports the equivalent results for the OrthoBench-calibrated simulations. OrthoFinder again had the highest overall composite score and the highest ortholog F1-score, and FastOMA the lowest of both. SonicParanoid2 and Broccoli had less orthogroup fission than OrthoFinder (Supplementary Figures S1.1 and S1.2).

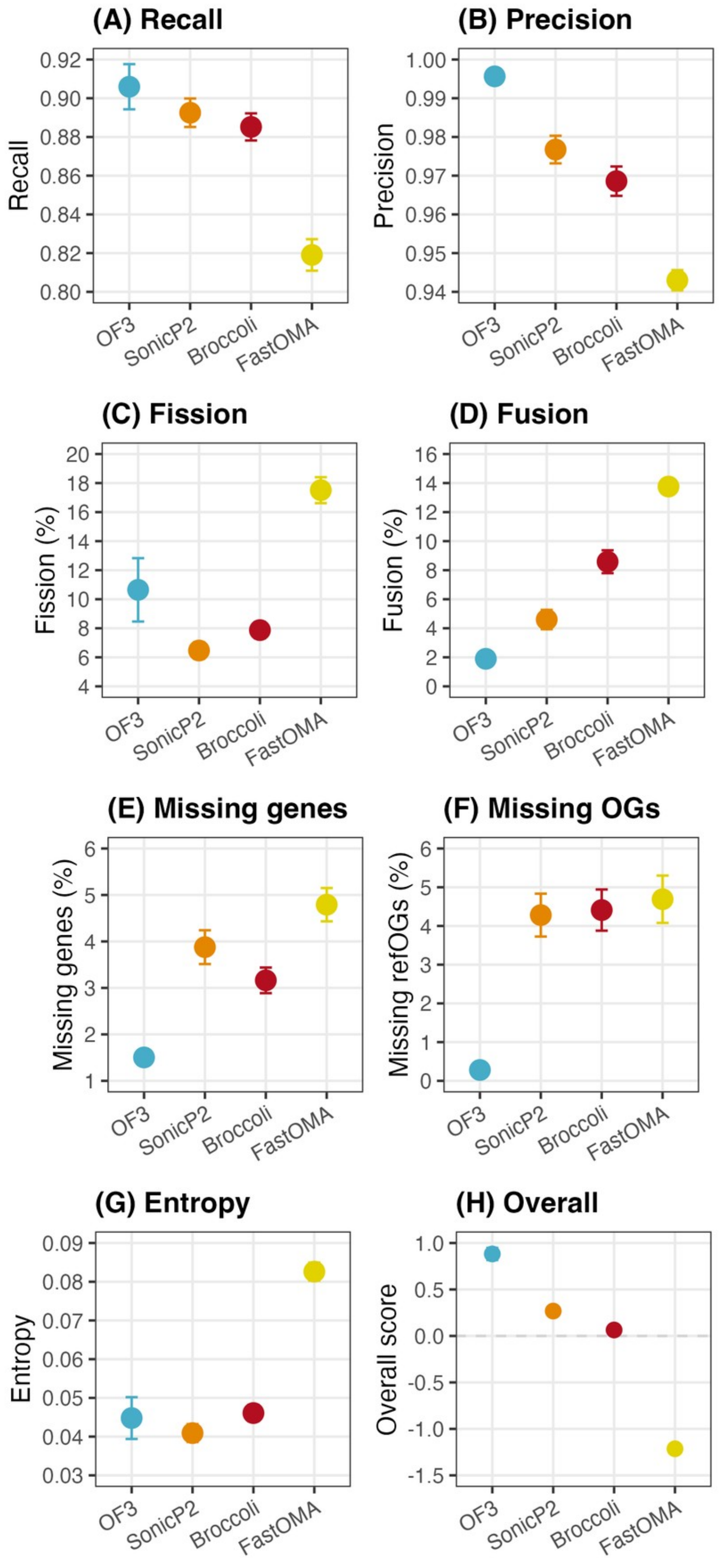


**Supplementary Figure S1.1** Orthogroup-level benchmarking of four orthology inference methods on the OrthoBench-calibrated simulations, equivalent to Figure 3. (A) Recall. (B) Precision. (C) Fission. (D) Fusion. (E) Missing genes. (F) Missing OGs. (G) Entropy. (H) Overall composite z-score.

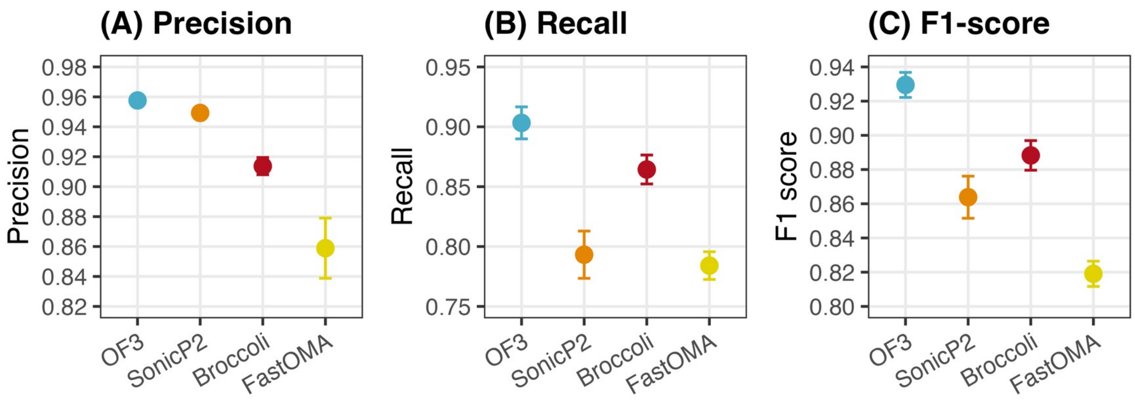


**Supplementary Figure S1.2** Ortholog-pair benchmarking of four orthology inference methods on the OrthoBench-calibrated simulations, equivalent to Figure 4. (A) Precision. (B) Recall. (C) F1-score.

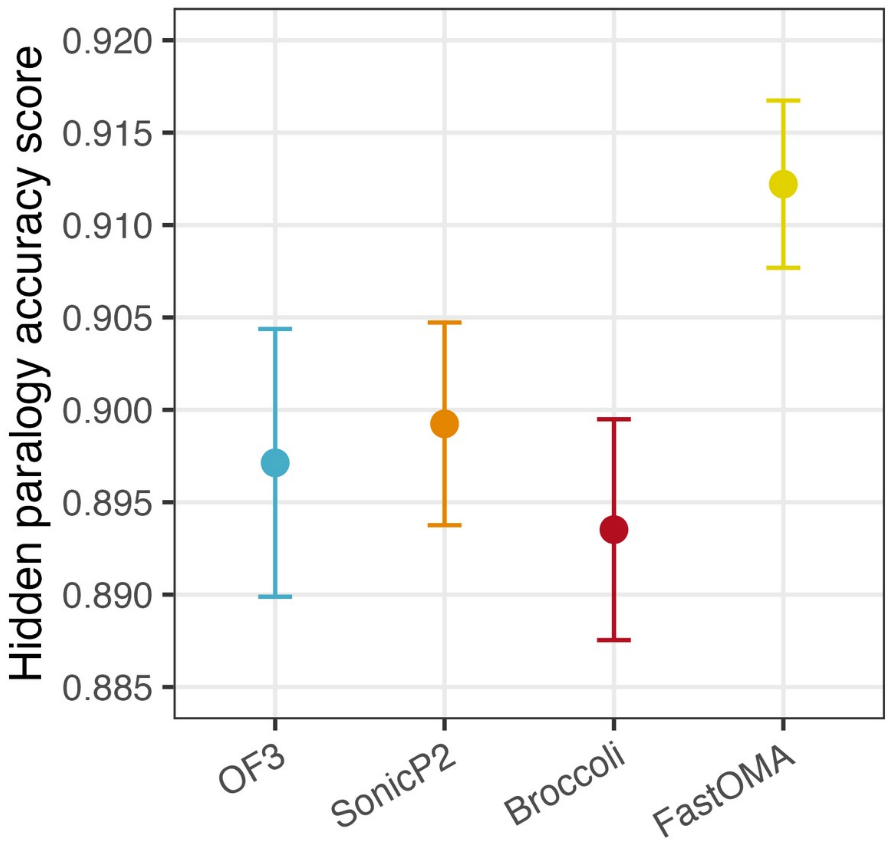


**Supplementary Figure S1.3** Hidden paralogy rejection rate for four orthology inference methods on the OrthoBench-calibrated simulations, equivalent to Figure 5.

#### Supplementary Figure 8

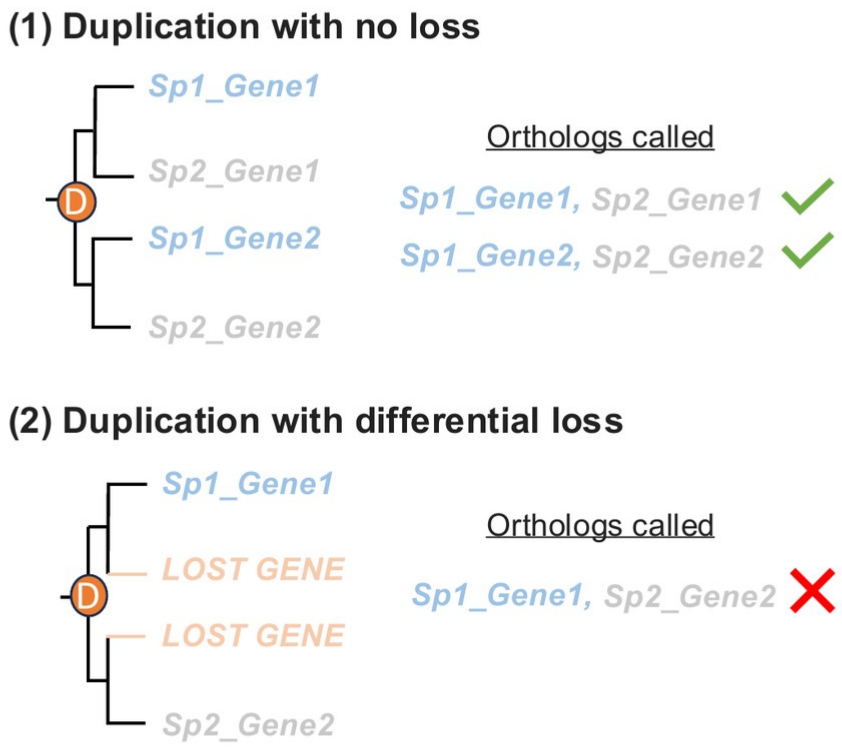


**Supplementary Figure 8**. Schematic illustration of hidden paralogy pairs. (1) A duplication with no subsequent loss. Both copies survive in both species, so orthologs are called correctly in both pairs. (2) A duplication followed by differential loss. Each species independently loses a different copy, leaving one gene per species that appear orthologous on a correct gene tree but are in fact paralogs descended through the duplication node, a hidden paralog pair.

#### Supplementary Figure 9

For each tool, we compared its performance on the simulation calibrated from its own empirical orthogroups against its mean performance on the simulations calibrated from the other three methods, after adjusting for the overall difficulty of each calibration dataset. Positive values indicate an advantage on own-calibrated data; the dashed line marks zero.

All four tools show small deltas throughout, indicating that the main-text rankings are not an artefact of which tool's orthogroups were used to calibrate the simulation.

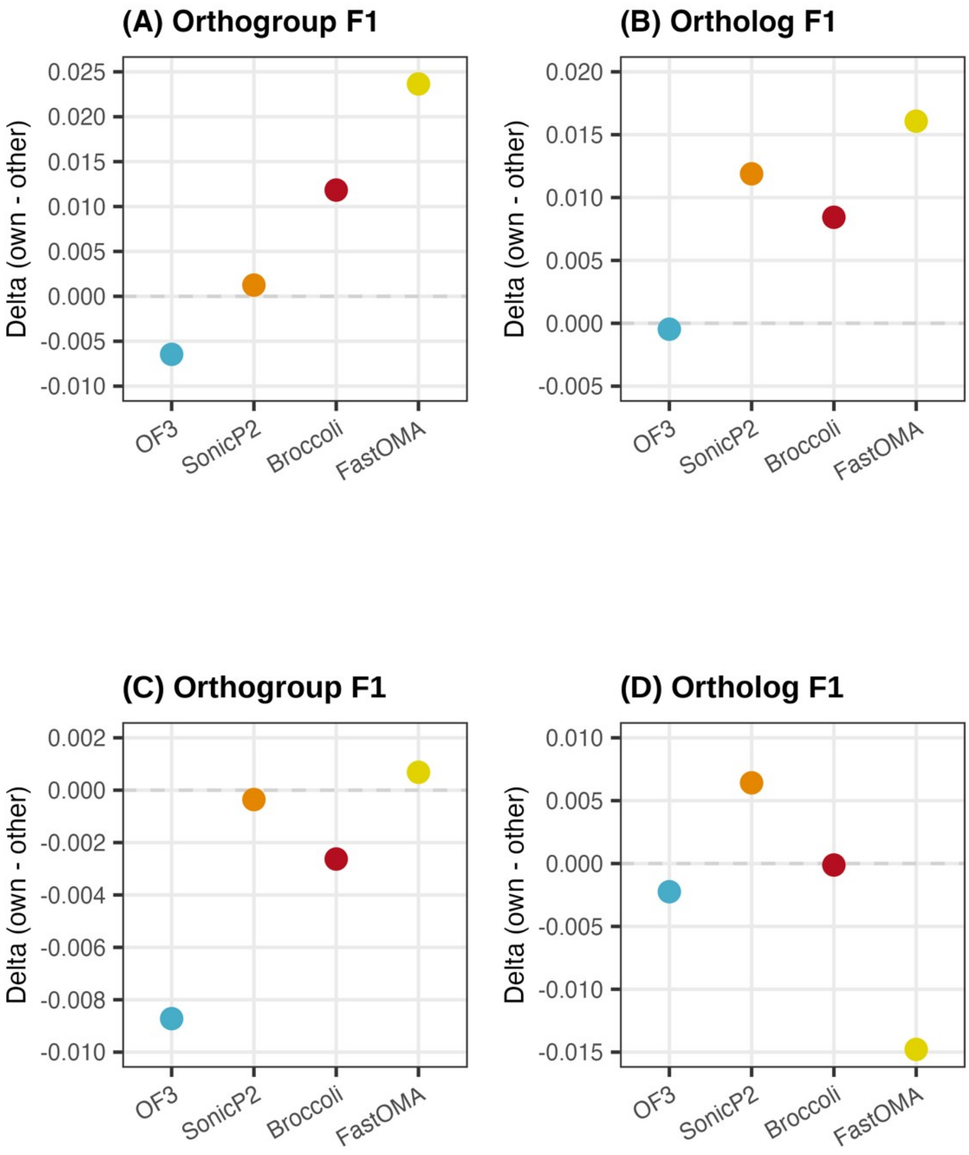


**Supplementary Figure 9**. Self-calibration advantage for orthogroup and ortholog inference. The top row is for the QfO dataset. The bottom row is for the OrthoBench dataset. (A&C) show Orthogroup F1, and (B&D) show Ortholog F1.

#### Supplementary Table 2

For each of the seven orthogroup- and ortholog-level benchmark measures reported in the main text, we compared the published OrthoBench ranking of the four orthology methods against the ranking recovered from our OrthoFinder-calibrated simulation. We drew 1,000 independent samples of 70 reference orthogroups, without replacement, from the full simulated reference set, recomputed all seven measures for each tool on each sample, and recorded, for every pair of tools, which one scored better. Where the published OrthoBench values for a pair of tools were themselves tied, three of six tool pairs on Missing RefOGs and RefOG Fusions, no comparison was made for that pair. This left 36 well-defined pairwise comparisons, shown in the table below, with Agreement (%) giving the fraction of the 1,000 resamples in which the simulation's pairwise comparison matched the published direction.

Agreement with the published direction was high overall (median 87% across the 36 comparisons, recomputed here as 86.55%), and comparisons involving FastOMA, the lowest-ranked tool in the published benchmark, were mostly reproduced with strong consistency (above 75% in 14 of 21 cases). A smaller number of comparisons (11 of 36) showed the opposite direction to the published result in a majority of resamples, and these were not evenly distributed. OrthoFinder versus SonicParanoid2 specifically recurred as a discordant result on three separate measures (RefOG Fissions, 1.6%; Precision, 5.6%; Entropy, 10.9%). One plausible contributor is that OrthoBench's reference set is deliberately enriched for orthogroups exemplifying known challenges for orthology inference rather than randomly sampled (Trachana et al. 2011), so two closely-matched tools' relative ranking on that enriched set need not match their ranking on a uniformly random sample of the kind this resampling draws. This is presented as a hypothesis rather than a conclusion; eight of the 11 involved SonicParanoid2, so the discordance centres on one tool rather than indicating a general mismatch between simulation and benchmark.

**Supplementary Table 2**. Bootstrap validation of OrthoBench-calibrated simulations against the published OrthoBench benchmark. This is the analysis underlying the main text's headline statistic of median 87% agreement across 36 pairwise comparisons.

| Metric | Tool A | Tool B | Published direction | Agreement (%) |
|---|---|---|---|---|
| Missing Refog | Broccoli | FastOMA | Broccoli better (published) | 72.0 |
| Missing Refog | FastOMA | OrthoFinder | OrthoFinder better (published) | 96.5 |
| Missing Refog | FastOMA | SonicParanoid2 | SonicParanoid2 better (published) | 44.9 |
| Missing Genes | Broccoli | FastOMA | Broccoli better (published) | 86.6 |
| Missing Genes | Broccoli | OrthoFinder | OrthoFinder better (published) | 98.4 |
| Missing Genes | Broccoli | SonicParanoid2 | Broccoli better (published) | 88.7 |
| Missing | FastOMA | OrthoFinder | OrthoFinder better | 100.0 |

| Metric | Tool A | Tool B | Published direction | Agreement (%) |
|---|---|---|---|---|
| Genes | | | (published) | |
| Missing Genes | FastOMA | SonicParanoid2 | SonicParanoid2 better (published) | 62.6 |
| Missing Genes | OrthoFinder | SonicParanoid2 | OrthoFinder better (published) | 100.0 |
| Fusion | Broccoli | FastOMA | Broccoli better (published) | 97.7 |
| Fusion | FastOMA | OrthoFinder | OrthoFinder better (published) | 23.4 |
| Fusion | FastOMA | SonicParanoid2 | SonicParanoid2 better (published) | 23.2 |
| Fission | Broccoli | FastOMA | Broccoli better (published) | 99.5 |
| Fission | Broccoli | OrthoFinder | Broccoli better (published) | 97.4 |
| Fission | Broccoli | SonicParanoid2 | Broccoli better (published) | 38.8 |
| Fission | FastOMA | OrthoFinder | OrthoFinder better (published) | 66.5 |
| Fission | FastOMA | SonicParanoid2 | SonicParanoid2 better (published) | 99.6 |
| Fission | OrthoFinder | SonicParanoid2 | OrthoFinder better (published) | 1.6 |
| Recall | Broccoli | FastOMA | Broccoli better (published) | 94.5 |
| Recall | Broccoli | OrthoFinder | OrthoFinder better (published) | 61.1 |
| Recall | Broccoli | SonicParanoid2 | Broccoli better (published) | 39.8 |
| Recall | FastOMA | OrthoFinder | OrthoFinder better (published) | 95.4 |
| Recall | FastOMA | SonicParanoid2 | SonicParanoid2 better (published) | 96.4 |
| Recall | OrthoFinder | SonicParanoid2 | OrthoFinder better (published) | 52.7 |
| Precision | Broccoli | FastOMA | FastOMA better (published) | 45.6 |
| Precision | Broccoli | OrthoFinder | OrthoFinder better (published) | 100.0 |
| Precision | Broccoli | SonicParanoid2 | SonicParanoid2 better (published) | 86.5 |

| Metric | Tool A | Tool B | Published direction | Agreement (%) |
|---|---|---|---|---|
| Precision | FastOMA | OrthoFinder | OrthoFinder better (published) | 98.9 |
| Precision | FastOMA | SonicParanoid2 | SonicParanoid2 better (published) | 79.0 |
| Precision | OrthoFinder | SonicParanoid2 | SonicParanoid2 better (published) | 5.6 |
| Entropy | Broccoli | FastOMA | Broccoli better (published) | 97.0 |
| Entropy | Broccoli | OrthoFinder | OrthoFinder better (published) | 28.9 |
| Entropy | Broccoli | SonicParanoid2 | Broccoli better (published) | 34.2 |
| Entropy | FastOMA | OrthoFinder | OrthoFinder better (published) | 89.9 |
| Entropy | FastOMA | SonicParanoid2 | SonicParanoid2 better (published) | 98.3 |
| Entropy | OrthoFinder | SonicParanoid2 | OrthoFinder better (published) | 10.9 |

**Secondary check: Spearman rank concordance**

As a complementary check, we also compared the four tools' OrthoBench-calibrated, OrthoBench-tested orthogroup benchmark scores, averaged across the four calibration datasets, against the published OrthoBench results for the same four tools and matching parameter settings (OrthoFinder, alignment-based species-tree mode; SonicParanoid2, default mode; Broccoli; FastOMA). All eight measures showed positive concordance (Spearman rho 0.40 to 1.00), with missing genes, F1, and recall showing the strongest agreement, rho at least 0.8.

As we only compare four tools, we analyse these data with an exact permutation test, computing the Spearman correlation for every one of the 24 possible tool relabelings and asking where the true correspondence ranked among them. Jointly across all eight measures, the true labeling achieved a higher mean concordance (rho = 0.69) than 22 of the 23 possible mismatched relabelings (p = 0.083), close to the best possible outcome given only four tools. Missing genes alone reached the single best possible result (rho = 1, p = 0.042). These results indicate that OrthoSimulator-derived rankings broadly agree with those obtained on the OrthoBench gold-standard benchmark, providing external support for OrthoSimulator's utility as a benchmarking resource, consistent with the bootstrap-resampling result above.

| **Metric** | **n** | **Spearman rho** | **p-value (exact, 24 relabelings)** |
|---|---|---|---|
| Missing genes | 4 | 1.00 | 0.042 |
| F1 | 4 | 0.80 | 0.167 |
| Recall | 4 | 0.80 | 0.167 |
| Missing RefOGs | 4 | 0.77 | 0.250 |
| Fusion | 4 | 0.77 | 0.250 |
| Precision | 4 | 0.60 | 0.208 |
| Fission | 4 | 0.40 | 0.375 |
| Entropy | 4 | 0.40 | 0.375 |
| Joint (all 8 metrics) | - | 0.69 | 0.083 |

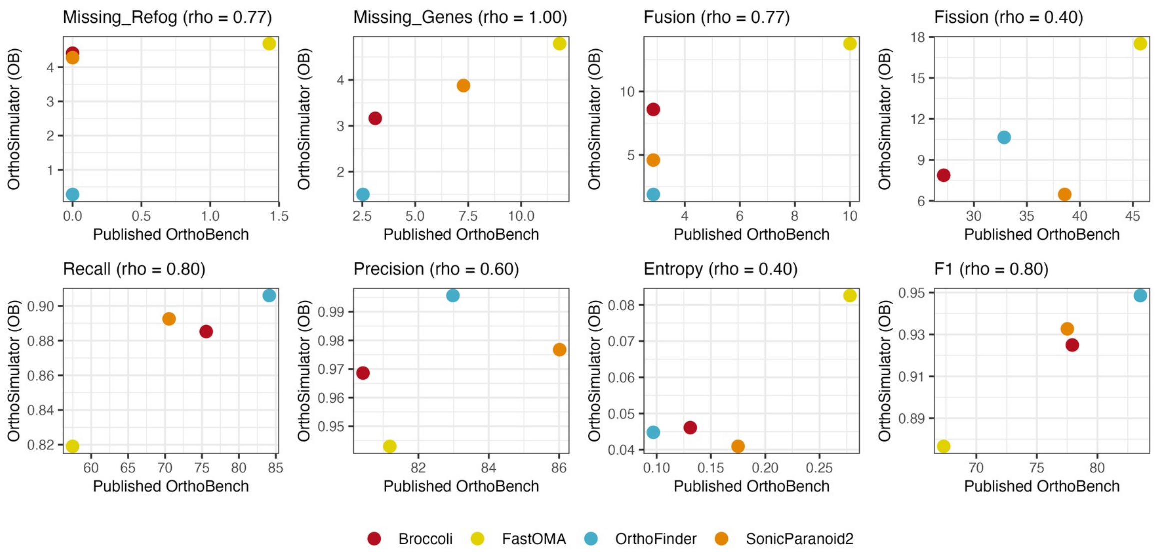


Per-metric concordance between OrthoBench-calibrated simulation results and the published OrthoBench benchmark. Each point is one of the four orthology inference tools; the diagonal indicates perfect rank agreement.

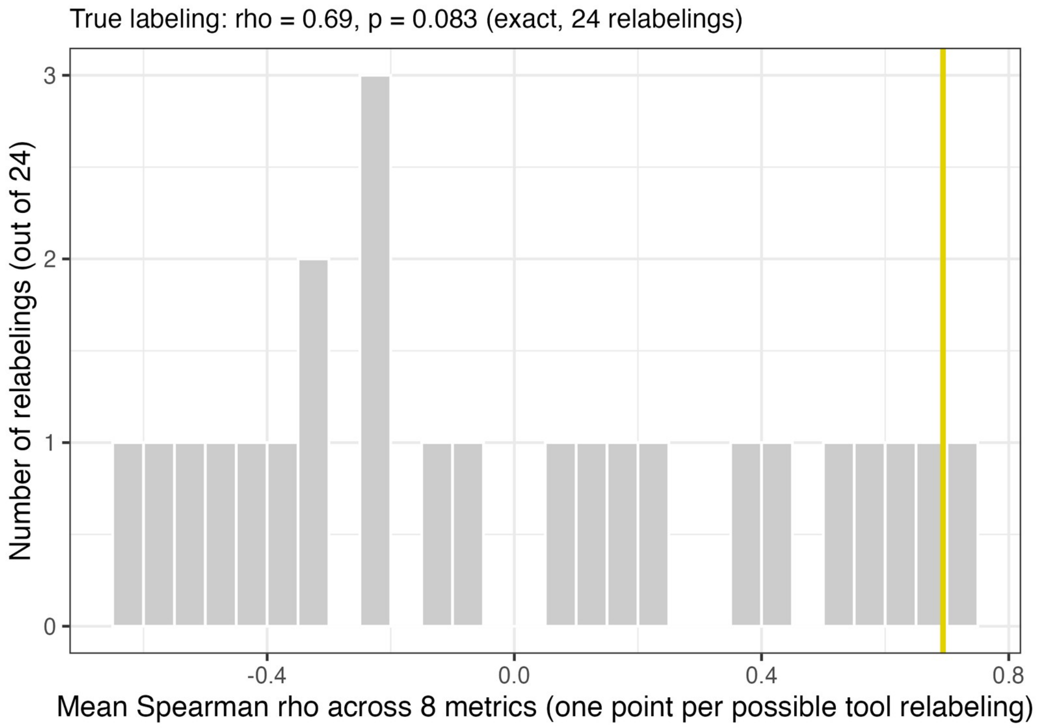


Permutation-test null distributions for the concordance statistics above. The observed, true-labeling Spearman rho is marked against the distribution of rho values from the 23 possible mismatched tool relabelings.

#### Supplement S2: Held-out validation details and figures

Several of the summary statistics in the main realism-validation figure are direct targets of the genetic algorithm's fitness function. To provide a more independent check of simulation realism, we compared simulated and empirical orthogroups on three further measures that are never used as simulation inputs or optimisation targets. These measures are (1) Sequence length. We measure this as mean ungapped sequence length per orthogroup. (2) Copy-number entropy. We measure this as the Shannon entropy of the per-species gene-copy-number vector for each orthogroup, including species with zero copies. (3) Pairwise sequence similarity. We measure this as the mean pairwise similarity across sequences in an orthogroup, scored under the JTT substitution matrix as the fraction of aligned residue positions with a positive substitution score.

The table below shows a quantitative summary of the held-out validation panel for the QfO dataset. W1 (sim, empirical) is the Wasserstein distance between simulated and empirical distributions for that measure. The final column expresses this as a ratio to the maximum of the six pairwise Wasserstein distances among the four empirical tools' own outputs on the same measure.

| **Calibrated from** | **Measure** | **W1 (sim, empirical)** | **Ratio to max empirical W1** |
|---|---|---|---|
| OrthoFinder | Sequence length | 59.96 | 0.822 |
| OrthoFinder | Copy-number entropy | 0.348 | 0.524 |
| OrthoFinder | Pairwise similarity (JTT) | 0.125 | 2.132 |
| FastOMA | Sequence length | 25.14 | 0.345 |
| FastOMA | Copy-number entropy | 0.342 | 0.514 |
| FastOMA | Pairwise similarity (JTT) | 0.090 | 1.530 |
| Broccoli | Sequence length | 45.57 | 0.625 |
| Broccoli | Copy-number entropy | 0.261 | 0.393 |
| Broccoli | Pairwise similarity (JTT) | 0.078 | 1.317 |
| SonicParanoid 2 | Sequence length | 15.88 | 0.218 |
| SonicParanoid 2 | Copy-number entropy | 0.442 | 0.665 |
| SonicParanoid 2 | Pairwise similarity (JTT) | 0.077 | 1.311 |

Sequence length and copy-number entropy fell within, or close to, the spread already observed among the four methods' own empirical distributions for all four methods (ratios 0.22 to 0.82 for sequence length; 0.39 to 0.67 for copy-number entropy). Pairwise similarity was the one measure where this did not hold. For every method, simulated sequences were consistently slightly more similar to one another than empirical sequences within the same orthogroup (ratios 1.31 to 2.13), possibly reflecting AliSim's sequence-generation mechanics rather than a mis-specified rate-heterogeneity model, since gamma shape and proportion invariant already match closely (Figure 2D, E).

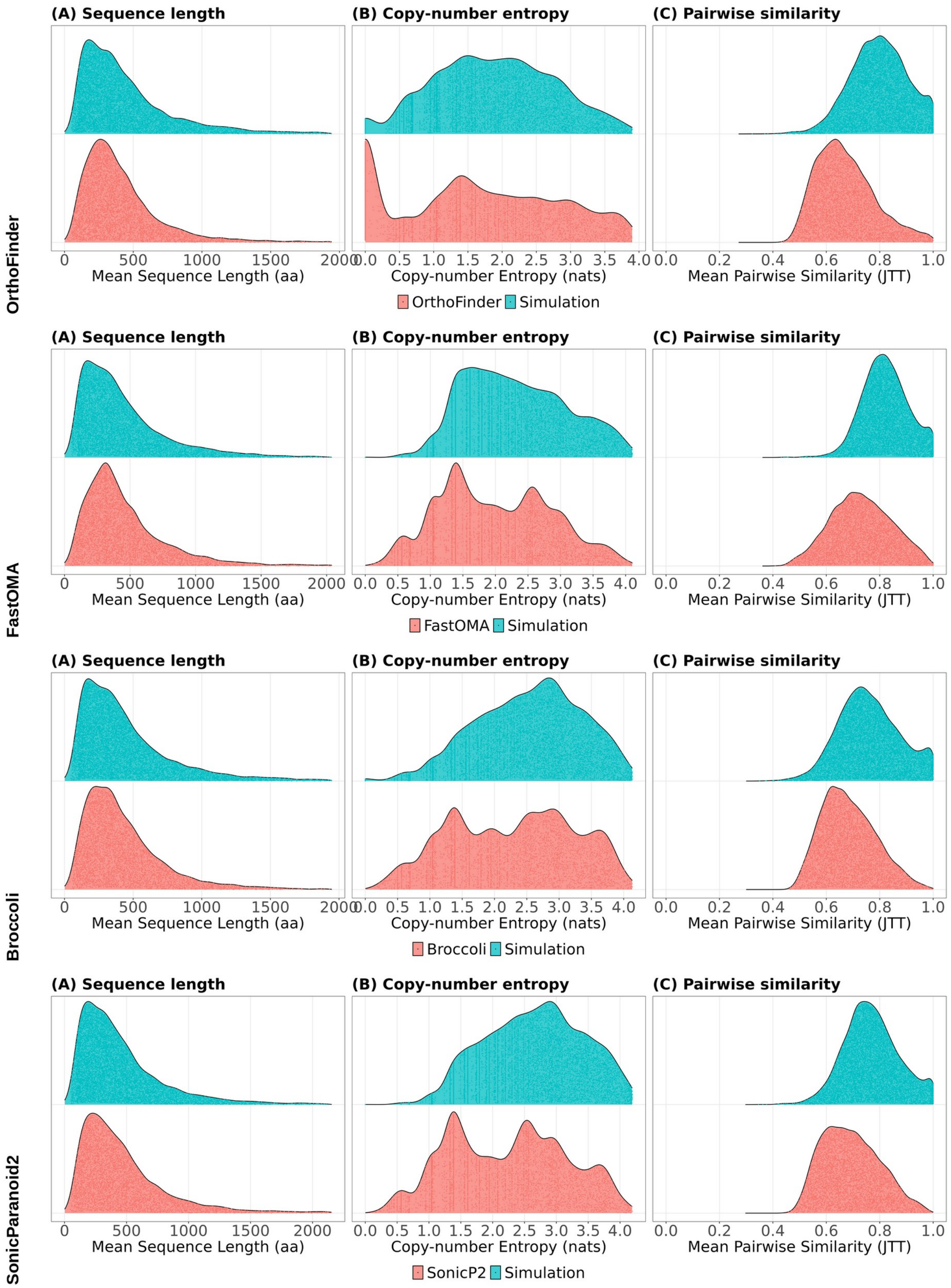


Held-out validation panel for the QfO dataset. Empirical (red) and simulated (blue) distributions for three measures not used in simulation calibration or parameter optimisation. Rows, top to bottom: OrthoFinder, FastOMA, Broccoli, SonicParanoid2. (A) Mean sequence length per orthogroup. (B) Copy-number entropy per orthogroup. (C) Mean pairwise sequence similarity per orthogroup, JTT substitution matrix.

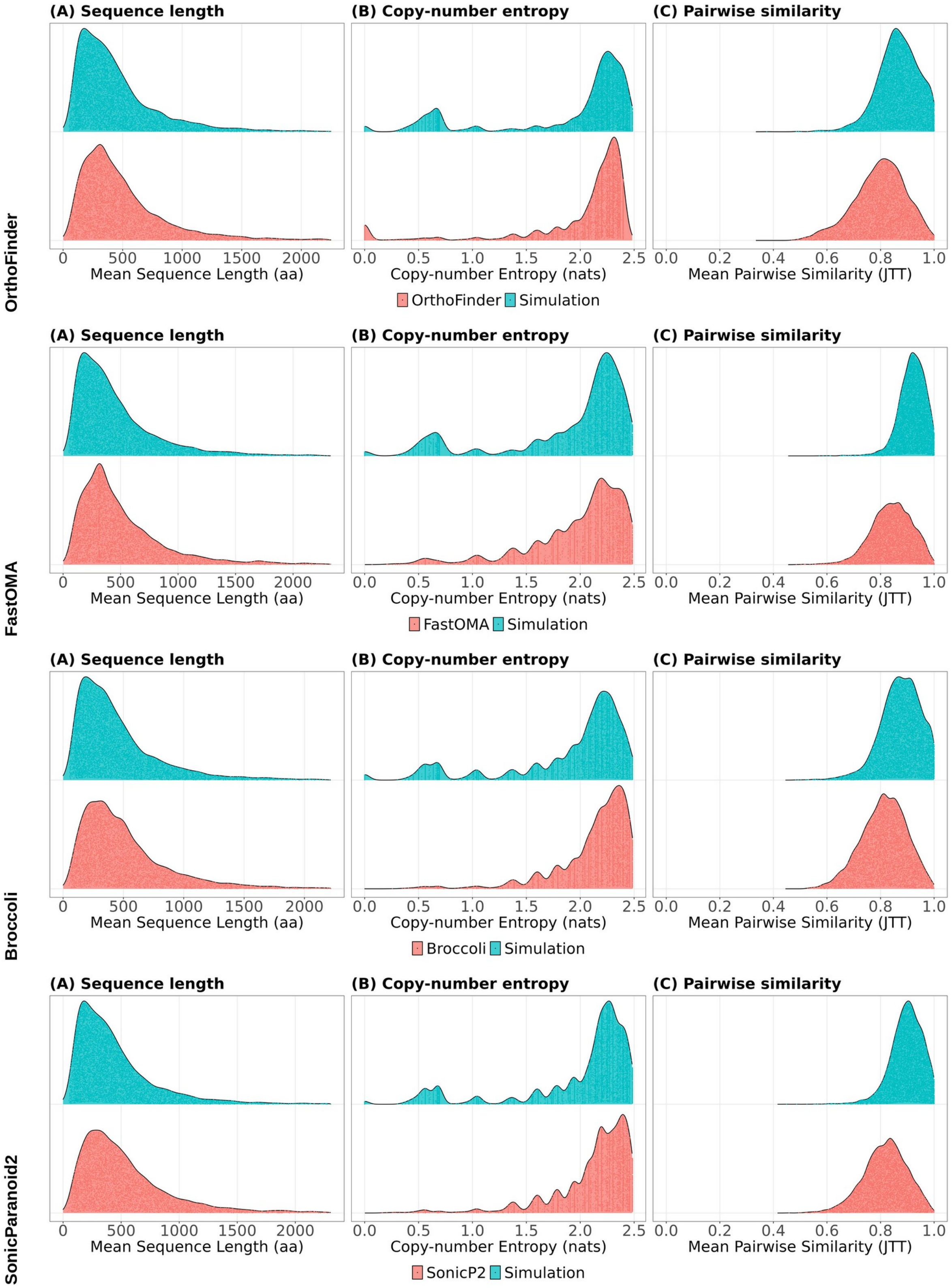


Held-out validation panel, OrthoBench dataset. As above, for the OrthoBench-calibrated simulations.

#### Supplementary Figure 10

SaGePhy (Kundu and Bansal 2019) simulates gene duplication, loss, and transfer under a birth-death process in which gene lineages coalesce at speciation. OrthoSimulator therefore uses this as its baseline gene-tree model, while also providing an optional extension that introduces incomplete lineage sorting (ILS) when this is required for a particular benchmarking question. Because the appropriate magnitude of ILS cannot generally be inferred from empirical orthogroup assignments alone, this extension is controlled by a single global parameter rather than being imposed by default.

The extension applies a post-hoc rearrangement step to each simulated gene tree so that its topology is consistent with the multispecies coalescent (Rannala and Yang 2003; Degnan and Salter 2005), while retaining duplication events as fixed, dated ground truth. Coalescence within each branch follows Kingman's coalescent. Species-tree branch lengths are converted to coalescent units using a single parameter, theta, defined such that branch length in coalescent units equals theta multiplied by branch length expressed as a fraction of root-to-tip height. Theta therefore controls the extent of ILS, with smaller values producing greater gene tree-species tree discordance and larger values producing less. Duplication events are retained using the bounded coalescent, so that gene copies descended from a duplication are constrained to have coalesced by the duplication time (Rasmussen and Kellis 2012).

To demonstrate this capability, we simulated an additional 10,000 orthogroups using the same calibrated parameters as the OrthoFinder-QfO baseline simulation, with theta = 10, and repeated the benchmarking of all four orthology inference methods. Quantitative results are summarised in the main text and the full benchmark results are shown below. This analysis demonstrates how OrthoSimulator can be used to extend ground-truth benchmarking to datasets containing explicitly simulated ILS and to examine how gene tree-species tree discordance affects orthology inference.

#### (A) Orthogroup-level benchmarking

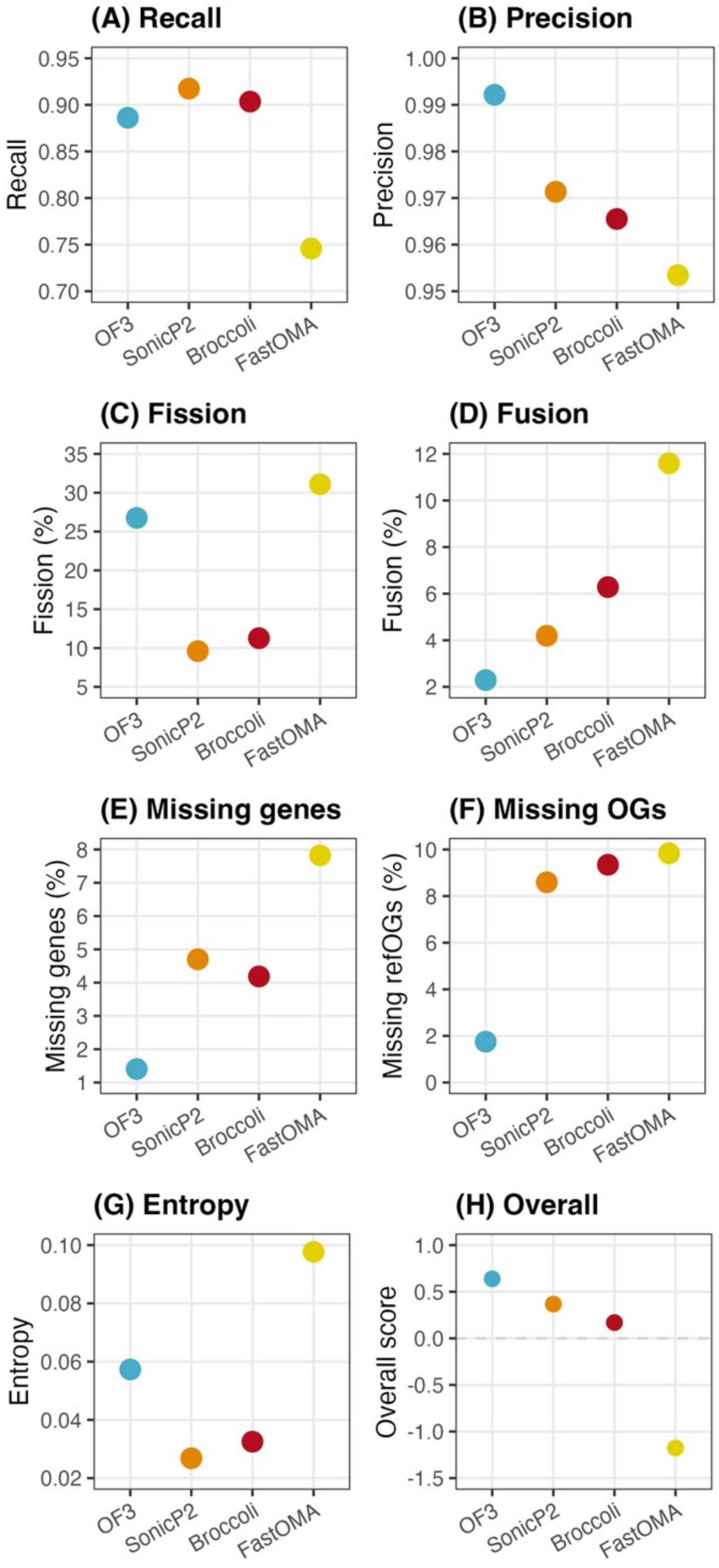

#### (B) Ortholog-pair benchmarking

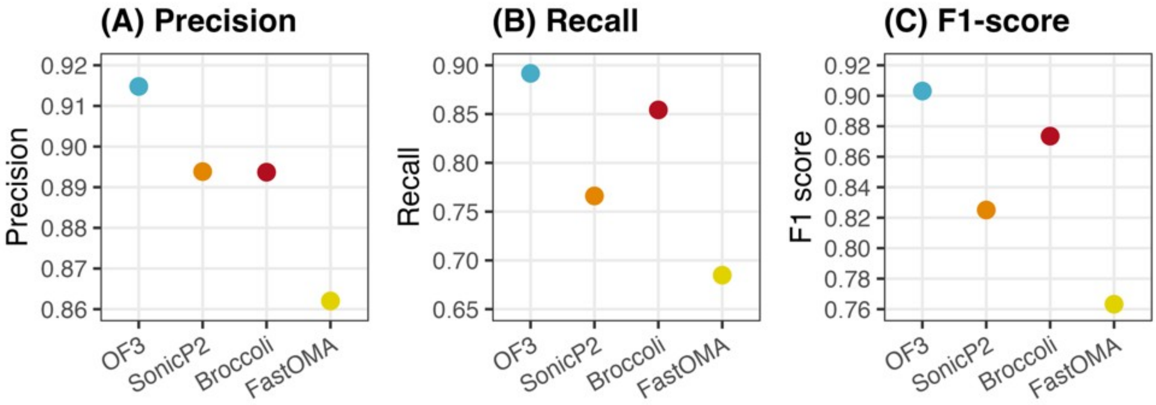


#### (C) Hidden-paralogy accuracy

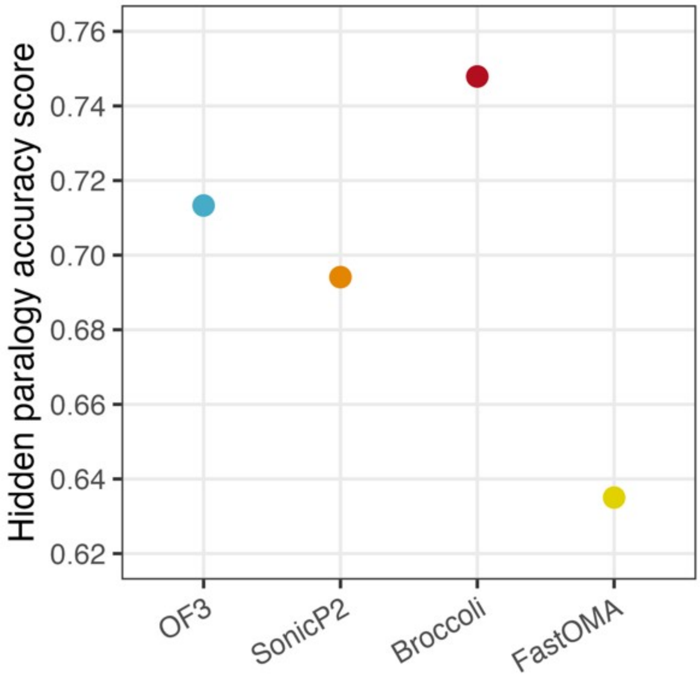


**Supplementary Figure 10**. Benchmarking of four orthology inference methods under simulated incomplete lineage sorting (theta = 10), QfO dataset, OrthoFinder-calibrated simulation. (A) Orthogroup-level benchmarking, equivalent to Figure 3. (B) Ortholog-pair benchmarking, equivalent to Figure 4. (C) Hidden-paralogy rejection rate, equivalent to Figure 5.

#### Simulation parameters

Three simulation parameters, the maximum duplication rate, the maximum loss rate, and the orthogroup birth-bias parameter gbc, cannot be reliably estimated directly from orthogroup assignments, because orthogroup-level data conflates rate and time and depends on the accuracy of the inference method used for empirical calibration. We therefore optimised these three parameters using a genetic algorithm that iteratively simulates small datasets and compares summary statistics of the simulated data to those of the empirical orthogroups (Methods, Genetic algorithm parameter optimisation). The table below reports the fitted value of each parameter for each of the four calibration methods, QfO dataset, together with their cross-tool mean, standard deviation, and coefficient of variation.

Supplementary Table 3. Genetic-algorithm-optimised simulation parameters and their variability across the four calibration methods, QfO dataset. gbc is the orthogroup birth-bias parameter. CV (%) is the coefficient of variation, SD divided by mean and multiplied by 100, of each fitted parameter across the four calibration methods.

| **Parameter** | **OrthoFinder** | **FastOMA** | **Broccoli** | **SonicParanoid2** | **Mean** | **SD** | **CV (%)** |
|---|---|---|---|---|---|---|---|
| gbc | 4.72 | 8.82 | 6.56 | 6.84 | 6.74 | 1.68 | 25.0 |
| max_duplication_rate | 3.93 | 1.04 | 1.75 | 0.67 | 1.85 | 1.46 | 79.1 |
| max_loss_rate | 2.34 | 5.09 | 2.14 | 2.05 | 2.91 | 1.46 | 50.2 |
